# GABA_A_R-PPT1 palmitoylation homeostasis controls synaptic transmission and circuitry oscillation in CLN1 disease

**DOI:** 10.1101/2024.03.26.585929

**Authors:** Jia Tong, Jingjing Gao, Yawei Qi, Ziyan Gao, Yang Liu, Qianqian Wang, Tiangang Yuan, Minglong Ren, Guixia Yang, Zhaoyue Li, Jin Li, Hongyuan Sun, Xing Zhao, Junmei Li, Yeung-Yeung Leung, Yonghui Mu, Jiamin Xu, Chengbiao Lu, Shiyong Peng, Lihao Ge

**Affiliations:** The Third Affiliated Hospital of Xinxiang Medical University, Xinxiang, He’nan, 453000, China; Institute of Psychiatry and Neuroscience, Xinxiang Medical University, Xinxiang, He’nan, 453003, China; Institute of Brain Functional Genomics, East China Normal University, Shanghai, 200333, China; He’nan International Joint Laboratory for Non-invasive Neural Modulation, Department of Physiology and Pathology, School of Basic Medical Science, Xinxiang Medical University, Xinxiang, He’nan, 453003, China; Division of Brain Sciences, Imperial College Faculty of Medicine, Du Cane Road, London W12 0NN, the United Kingdom; Basic Medical College, Xinxiang Medical University, He’nan, 453003, China

**Keywords:** CLN1 disease, palmitoyl protein thioesterase 1, PPT1-KI, GABA_A_R, *in vivo* field potential (FP) recordings, synaptic transmission, higher brain function

## Abstract

CLN1 disease, also called infantile neuronal ceroid lipofuscinosis, is a fatal neurodegenerative disease caused by mutations in the CLN1 gene encoding palmitoyl protein thioesterase 1 (PPT1). To identify depalmitoylation substrate of PPT1 is crucial to understand CLN1 disease. In this study, we found that PPT1 depalmitoylates GABA_A_R α1 subunit at Cystein-260, while binding to Cystein-165 and −179. Mutations of PPT1 or its GABA_A_R α1 subunit binding site result in enhanced inhibitory synaptic transmission, strengthened and oscillation but disrupted phase coupling in CA1 region and impaired learning and memory in 1- to 2-months-old PPT1-deficient and *Gabra1^em1^* mice. Our study highlights the critical role of PPT1 in maintaining GABA_A_R palmitoylation homeostasis and reveals a previously unknown molecular pathway in PPT1 induced diseases.

## Introduction

Palmitoylation is a posttranslational modification in which a long-chain 16-carbon fatty acid (palmitate) is attached to the thiol groups of cysteine residues in substrate proteins (Camp et al., 1994). This modification influences the cellular distribution of substrate proteins and regulates protein function (Jin et al., 2021). The enzymes responsible for this process are palmitoyl acyltransferases, that contain a zinc-finger domain with a conserved DHHC (Asp-His-His-Cys) motif (zDHHCs) (Lemonidis et al., 2015; Mitchell et al., 2006). Depalmitoylation is a reverse process of palmitoylation and is catalysed by palmitoyl-protein thioesterases (PPTs), acyl-protein thioesterases, and the α/β hydrolase domain-containing proteins (Lin and Conibear, 2015b; Vesa et al., 1995).

Neuronal ceroid lipofuscinoses (NCLs) are worldwidely distributed rare diseases. Epidemiological studies indicate an incidence of 1–3/100,000 and a prevalence of about 2– 4/1,000,000(Santorelli et al., 2013; Sleat et al., 2016; Williams, 2011). Different forms of NCL are typically classified according to the age of onset, with infantile NCL (INCL) exhibiting the earliest onset and the most rapid course(Santavuori et al., 1973). Previous studies indicate that 25 % of the patients worldwide and 40 % in the United States harbour nonsense mutations in the *CLN1* encoding PPT1 gene (Bouchelion et al., 2014). Patients with CLN1 disease undergo progressive retinal degeneration, leading to blindness, learning and memory loss, and seizures(Hobert and Dawson, 2006; Mukherjee et al., 2019).

Despite the severity of PPT1 mutations during early brain development, the mechanisms underlying the PPT1 deficiency-induced clinical profile of CLN1 disease remain poorly understood. A large-scale proteomic screen has shown that more than 100 synaptic proteins are PPT1 substrates(Gorenberg et al., 2022), which depicts the complexity of PPT1 function in the central nervous system. Other studies have indicated that PPT1 plays important roles in synapses, such as regulating axonal growth(Chu-LaGraff et al., 2010) and neurite extension(Lange et al., 2018; Sapir et al., 2019). In the presynaptic area, PPT1 modifies the synaptic protein synaptobrevin 2 (VAMP2), the synaptosomal-associated protein of 25kD (SNAP 25), and the cysteine string protein α (CSPα), thereby PPT1 deficiency causes vesicle recycling failure in both mice and humans(Sung-Jo Kim). In the postsynaptic area, GluN2B and Fyn kinases are hyperpalmitoylated in PPT1^-/-^ neurons, which hinder the developmental *N*-methyl-D-aspartate receptor (NMDAR) subunit switch from GluN2B to GluN2A(Koster et al., 2019). Studies have shown that PSD-95 and GluN2A are not PPT1 substrates (Koster *et al*., 2019; Yokoi et al., 2016). To better understand the pathogenesis of CLN1 disease, we try to identify the synaptic depalmitoylation substrates of PPT1.

Further, GABA_A_R plays a central role in synchronising network neuronal activity, which are tightly associated with higher brain functions such as learning and memory(Nakazono et al., 2018). PPT1-KI mice carrying nonsense mutations in the *CLN1/PPT1* gene exhibited reduced expression of GABA_A_R along with impaired gamma oscillation and seizure onset (Zhang et al., 2022). These findings indicate the importance of GABA_A_R in CLN1 disease.

In the present work, we undertook multilevel functional and biochemical approaches to investigate both inhibitory and excitatory synaptic transmission, and generated transgenic mice *Gabra1^em1^* with a double mutation of GABA_A_R α1 binding sites to PPT1, aiming to elucidate the postsynaptic mechanisms of NCL. We demonstrated that the GABA_A_R α1 subunit is a substrate of PPT1. In 1- to 2-month-old PPT1-deficient (PPT1-KI) mice, we found inhibitory synaptic transmission was enhanced, GABA_A_R was hyperpalmitoylated, which resulted in membrane retention of GABA_A_R. *Gabra1^em1^* mice show disrupted neural network oscillations similar to those observed in PPT1-deficient mice. These results indicated that GABA_A_R, as a PPT1 substrate, via alteration of its palmitoylation which affect function of neural circuits in the early stage and provides a novel synaptic mechanism for NCL pathogenesis.

## Results

### PPT1-KI enhances GABA_A_R α1 membrane expression and GABAergic neurotransmission

To investigate the function of postsynaptic receptors in PPT1-deficiency induced diseases, we first assessed the role of GABA_A_R α1 subunit hyperpalmitoylation in regulating inhibitory neurotransmission. Consequently, we investigated GABAergic neurotransmission in 1- to 2-month-old PPT1-KI mice to simulate the early stages of pathogenesis of CLN1 disease. Evoked inhibitory postsynaptic currents (eIPSCs) were recorded by holding pyramidal cells in the CA1 region at +40 mV, as shown in Fig. 1A. Compared with wildtype (WT) neurons, PPT1-KI neurons showed an increased amplitude (Fig. 1B) and a decreased fast tau value without affecting the slow tau by double exponential fitting of the repolarization phase (Fig. 1C). Miniature inhibitory postsynaptic currents (mIPSCs) amplitude and frequency were both enhanced in PPT1-KI neurons, which is similar to *Gabra1^em1^*mice (Fig. 1D-F), as described below. Next, we incubated hippocampal slices of PPT1-KI mice with 1 μM BuHA, a thioesterase mimetic that selectively cleaves thioester linkage in palmitoylated proteins and compensates for the molecular defect caused by PPT1 mutations (Sarkar et al., 2013), both the amplitude and frequency of mIPSCs were recovered to the control level (Fig. 1D-F). These results demonstrated that hyperpalmitoylation of the GABA_A_R α1 subunit by PPT1 deficiency strengthened GABAergic neurotransmission.

**Figure 1.**
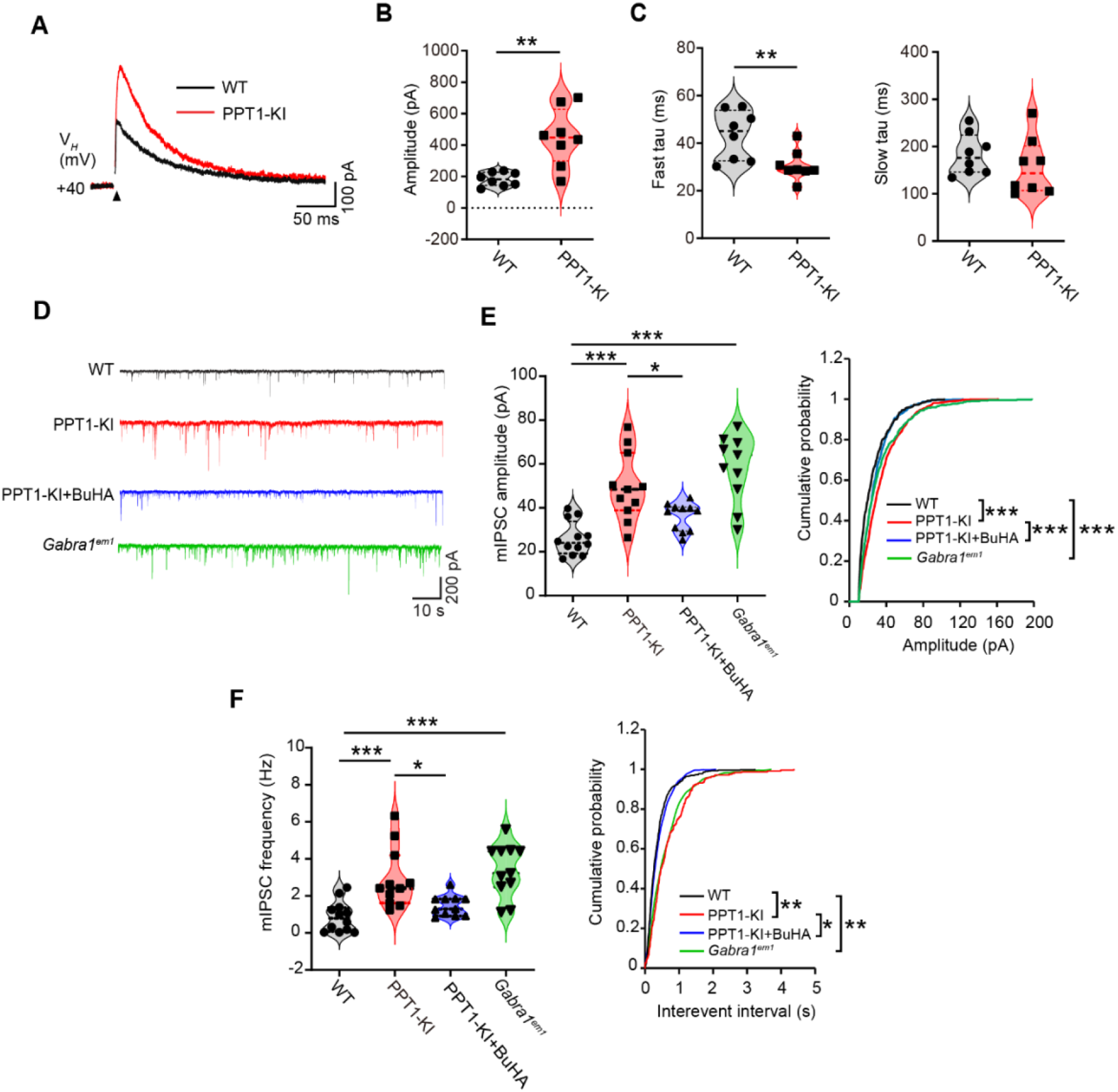
Mobilization of GABA_A_R α1 subunit to the cellular membrane enhances GABAergic transmission in PPT1-deficient mice. **(A)** Sample traces showing evoked IPSCs recorded at +40 mV from CA1 pyramidal cells in 1- to 2-month-old WT (black trace) and PPT1-KI mice (red trace). **(B, C)** Amplitude **(B)**, fast tau, and slow tau **(C)** of IPSCs from **A**. WT: n=8 neurons from six slices; PPT1-KI: n=8 neurons from five slices. T-test, \*\**P* < 0.01. **(D-F)** Sample traces **(D)** showing mIPSCs amplitude **(E)** and frequency **(F)** are enhanced in PPT1-KI mice, *Gabra1^em1^* mice, compared to WT mice, and partially recovered by pre-incubation with 1 μM BuHA. WT: n=12 neurons from 9 slices; PPT1-KI: n=11 neurons from 7 slices; *Gabra1^em1^*: n=11 neurons from 10 slices, PPT1-KI with BuHA: n=11 neurons from 6 slices. Histogram: One-way ANOVA, \**P* < 0.05, \*\**P* < 0.01, \*\*\**P* < 0.001; cumulative curve: Kolmogorov–Smirnov test, \*\**P* < 0.01, \*\*\**P* < 0.001. Data are represented as mean ± SEM.

Furthermore, we examined whether postsynaptic glutamatergic receptors could be affected by PPT1; AMPAR-and NMDAR-mediated currents (Fig. 2A-C), as well as the AMPA/NMDA current ratio (Fig. 2D) showed no difference between WT and PPT1 mice, miniature excitatory postsynaptic currents (mEPSCs) amplitude and frequency were also comparable in WT and PPT1-KI mice with and without BuHA treatment (Fig. 2E-G). Western blot analysis of AMPAR, NMDAR 2a/2b, and their scaffold proteins (PSD95/93 and SAP102) showed no changes in membrane protein expression in WT and PPT1-KI mice with and without BuHA treatment (Fig. S1). These data suggest that PPT1-KI did not affect glutamatergic transmission.

**Figure 2.**
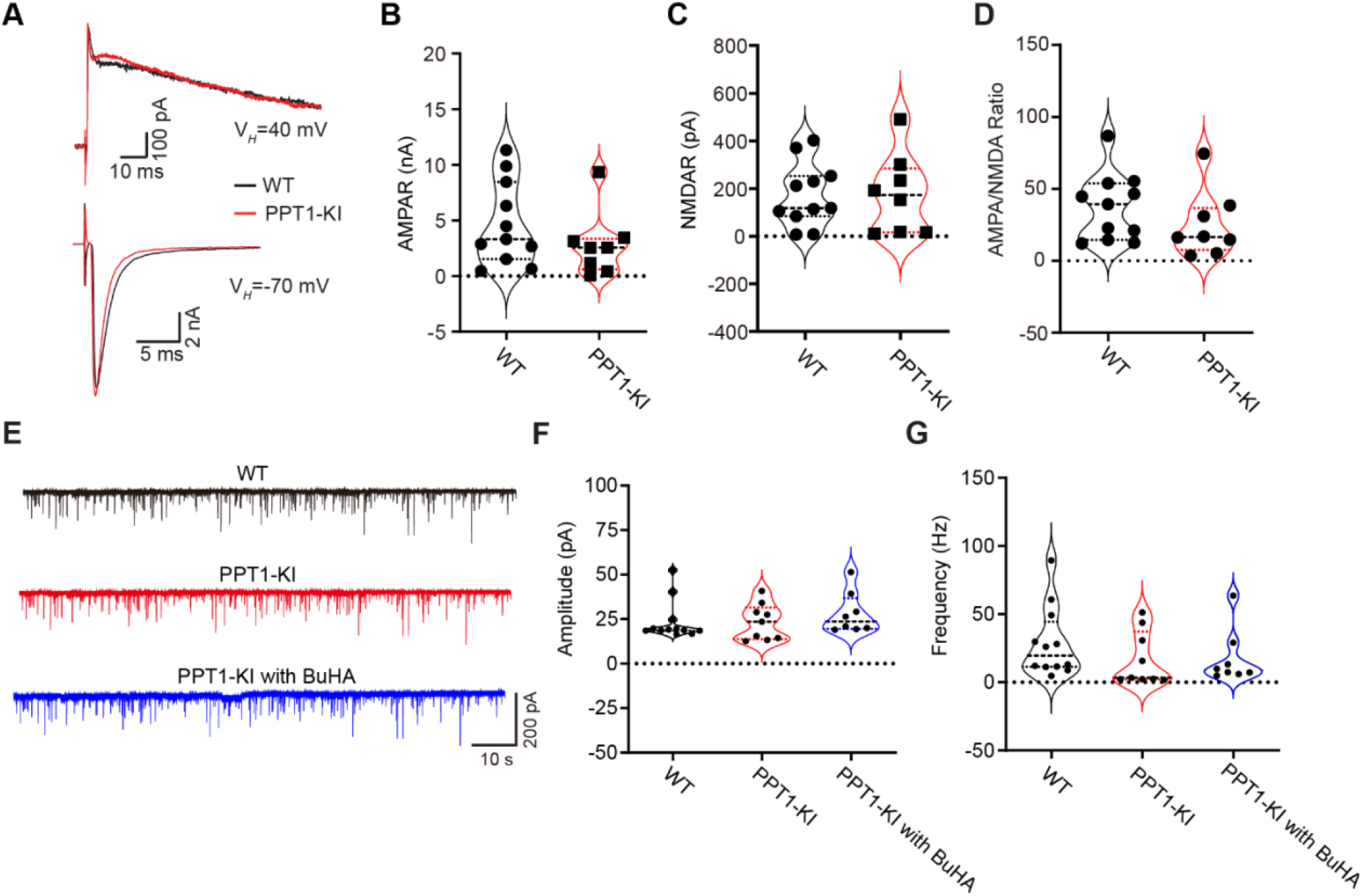
Excitatory synaptic transmission is not altered in PPT1-deficient mice. (**A**) Sample traces of evoked EPSCs recorded from WT (black trace) and PPT1-KI mice (red trace) hippocampal pyramidal neurons at +40 mV (upper traces) and −70 mV (bottom traces). (B-D) Amplitudes of AMPAR (**B**), NMDAR (**C**), and AMPAR/NMDA ratio (**D**) from the evoked EPSCs of hippocampal pyramidal neurons. WT: n=11 neurons from 10 slices; PPT1-KI: n=8 neurons from 6 slices; t-test: no significant difference. (**E-G**) Representative traces and analyses of mEPSCs recorded from CA1 pyramidal neurons of WT (black trace) and PPT1-KI (red trace) mice. WT: n=12 neurons from 10 slices; PPT1-KI: n=9 neurons from 9 slices; PPT1-KI with 5 μM BuHA: n=8 neurons from 7 slices. A one-way ANOVA showed no significant differences. Data are represented as mean ± SEM.

### GABA_A_R α1 subunit membrane expression is developmentally regulated in PPT1-KI mice

We tested the membrane expression of GABA_A_R α1, β2, and γ2 subunits at different developmental stages by performing membrane fraction analysis using western blot. In PPT1-KI mice, we observed a distinct age-dependent pattern for the α1 subunit: membrane expression was increased at 1 and 2 months but decreased at 7 months (Fig. 3A, B). While, qPCR results showed no differences in the total mRNA expression of all GABA_A_R subunits until 7 months (Fig. 3C). Furthermore, the membrane expression level of GABA_A_R α1 in PPT1-KI mice was higher than that of the WT, which could be mitigated by *in vitro* incubation with different concentrations of BuHA (Fig. 3D-E). The membrane expression of β2 and γ2 subunits remained unchanged compared to the WT group (Fig. S2). These results suggest that the increased membrane level of the α1 subunit in 1- to 2-month-old PPT1-KI mice is initiated by post-translational modification rather than changes in gene expression. Further, the reduced membrane level of the α1 subunit in 7-month-old mice is likely due to neuron loss(Kielar et al., 2007).

**Figure 3.**
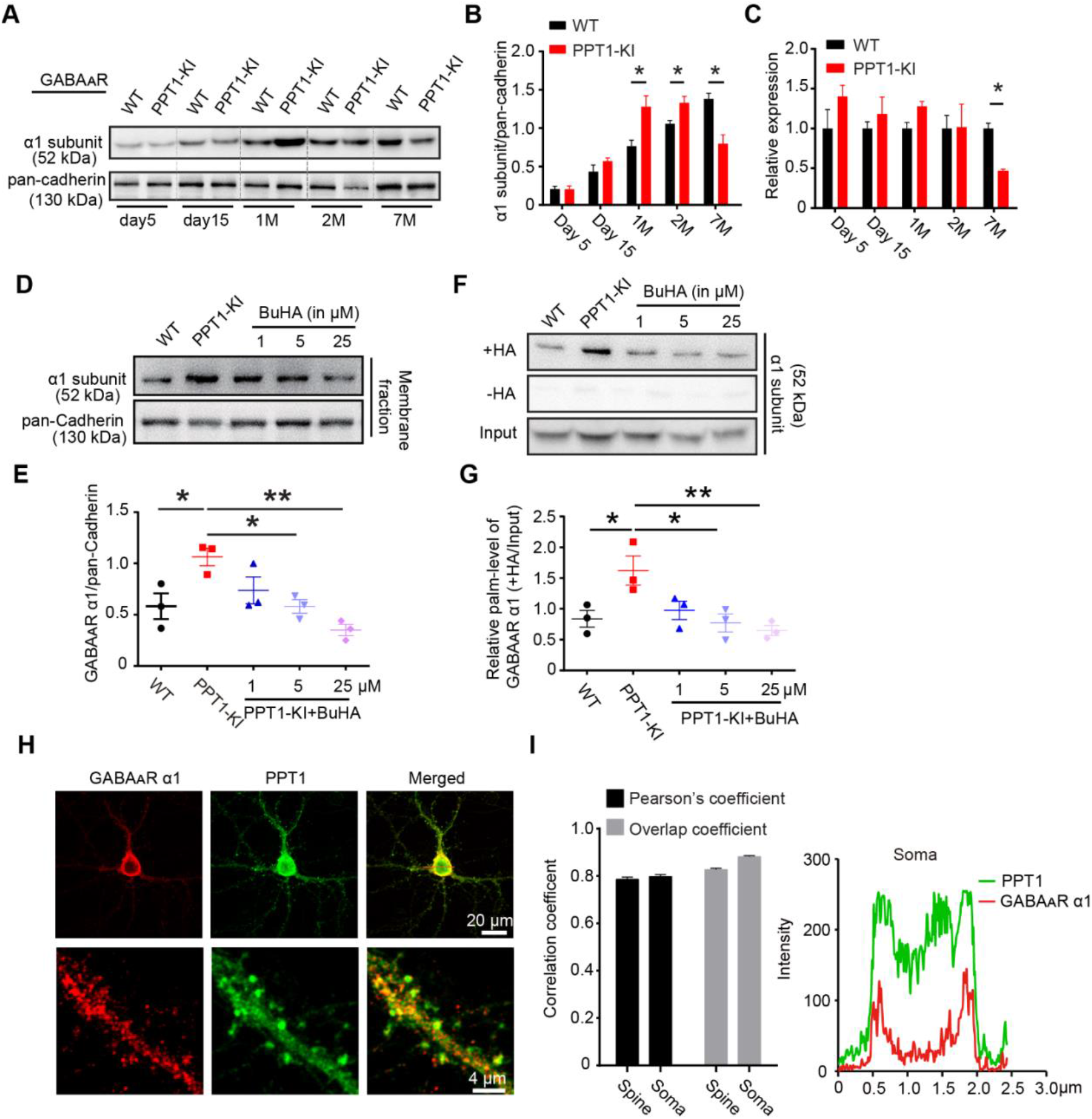
GABA_A_R α1 subunit is a substrate of PPT1 enzyme. **(A, B)** Representative immunoblot **(A)** and quantification **(B)** of GABA_A_R α1 subunits levels on the cellular membrane in WT and PPT1-KI mice at indicated ages. N=3 for each group, two-way ANOVA, \**P* < 0.05. **(C)** Quantitative PCR showing developmental changes in the mRNA expression of GABA_A_R α1 subunit at the indicated ages. N=3, two-way ANOVA, \**P* < 0.05. **(D, E)** Representative immunoblotting **(D)** and quantification **(E)** of GABA_A_R α1 subunit levels on the cellular membrane in WT, PPT1-KI, and PPT1-KI mice treated with BuHA. N=3 for each group, one-way ANOVA; \**P* < 0.05, \*\**P* < 0.01. **(F, G)** Increased levels of the palmitoylated GABA_A_R α1 subunit in cultured PPT1-KI neurons, which were recovered by incubation with BuHA. **(H, I)** Representative images **(H)** and colocalization analysis **(I)** of GABA_A_R α1 subunit and PPT1 enzyme in cultured hippocampal neurons from WT mice. Fluorescence intensity was measured using the red line across the cell body. N=8 cells. +HA, with hydroxylamine; N= 3 for each group, one-way ANOVA, \**P* < 0.05; \*\**P* < 0.01. Data are represented as mean ± SEM.

### PPT1 depalmitoylates GABA_A_R α1 subunit at cysteine residues

We performed an immunocytochemical assay to examine the subcellular distributions of the GABA_A_R α1 subunit and the PPT1 enzyme in hippocampal neurons. We found that the endogenous GABA_A_R α1 subunit colocalized with PPT1 in neuronal soma, neurites and spines with super-resolution stimulated emission depletion (STED) microscopy (neurites and spines: Pearson’s coefficient: 0.79± 0.0091, overlap coefficient: 0.83± 0.0060; soma: Pearson’s coefficient: 0.80± 0.0077, overlap coefficient: 0.88± 0.0041, n=8 cells) (Fig. 3H, I). The close localisation of the GABA_A_R α1 subunit and PPT1 prompts the investigation of whether the GABA_A_R α1 subunit can be palmitoylated by PPT1. We employed the Acyl-Biotin Exchange (ABE) assay to analyse the palmitoylation level of membrane fractions; the GABA_A_R α1 subunit’s palmitoylation level is upregulated in PPT1-KI mice, which was suppressed by *in vitro* incubation with BuHA (Fig. 3F, G). In contrast, the palmitoylation status of its postsynaptic scaffold protein gephyrin was not affected (Fig. S3), suggesting that an enzyme other than PPT1 regulates the depalmitoylation of gephyrin.

Moreover, an *in vitro* co-immunoprecipitation (co-IP) assay revealed the biochemical interaction between PPT1 and the GABA_A_R α1 subunit (Fig. 4A), indicating that GABA_A_R is a substrate of PPT1. Bioinformatics analysis predicted four potential palmitoylated cysteine residues (C165, C179, C260, and C319) (Fig. 4B). The mutation of GABA_A_R α1 subunit 260-cysteine residue to alanine reduces its palmitoylation level, suggesting the palmitoylation site (Fig. 4C and D). To identify the binding site between PPT1 and GABA_A_R α1, we mutated different GABA_A_R α1 cysteine residues to alanine in the 293T cell line. Co-IP results showed that single mutation of the cysteine residues did not affect PPT1-GABA_A_R α1 binding; however, this binding was strongly reduced when we double-mutated 165 and 179 cysteines to alanine (C165A& C179A) (Fig. 4E and F). In contrast, no reduction was detected when cysteine residues (260 and 319) were mutated to alanine residues (C260A and C319A) (Fig. 4G and H). These results indicate that C165 and C179 are the binding sites of GABA_A_R α1 to PPT1.

**Figure 4.**
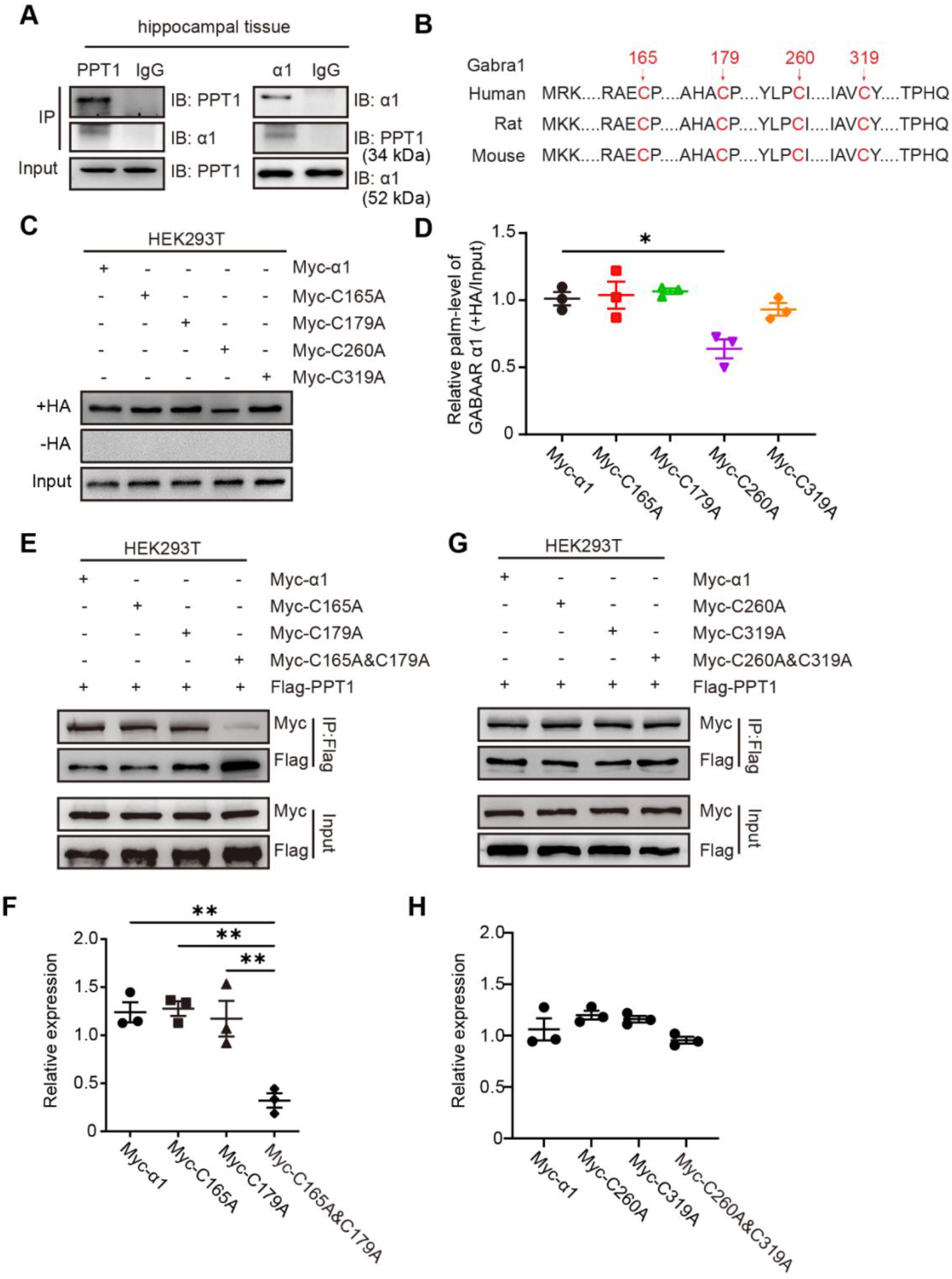
Identification of the binding sites between GABA_A_R α1 subunit and PPT1. **(A)** Interaction between PPT1 and GABA_A_R α1 subunit verified by *in vitro* CO-IP using mouse hippocampal tissue. IB: immunoblotting. **(B)** Bioinformatic prediction of palmitoylation site (Cys). (**C, D**) Identification of palmitoylation sites within GABA_A_R α1 subunit. Mutation of Cys-260 residues to alanine reduce palmitoylation level of GABA_A_R α1 in 293T cell line. N= 3, one-way ANOVA, \**P*< 0.05. **(E-H)** Representative blots and quantification of immunoprecipitation show that mutations of Cys-165 and Cys-179 **(E, F)**, but not Cys-260 or Cys-319 **(G, H)**, to alanine block GABA_A_R α1 subunit binding to PPT1 in 293T cell line. N= 3, one-way ANOVA, \*\**P*< 0.01.

We also investigated the potential of PPT1 as a substrate for excitatory synapses. The ABE results showed that the palmitoylation status of ionotropic glutamate receptors (iGluRs), such as AMPAR and NMDA 2a/2b, as well as their scaffold proteins PSD95/93 and SAP102, were comparable between PPT1-KI and WT mice (Fig. S4A–G). QPCR analysis showed no changes in mRNA expression at an early stage (Fig. S4H). Our data suggest that these excitatory synaptic proteins are unlikely to be PPT1 substrates.

### Disruption of PPT1 or GABA_A_R α1 subunit impairs neural network oscillations

A previous study reported that *CLN3^-/-^* mice, as young as 2 months old, exhibited a widespread decrease in network coordination in most of the hippocampus and fast beta frequency power with decreased slow delta activity by 18 months of age (Ahrens-Nicklas et al., 2019). Using *in vivo* field potential (FP) recordings of the hippocampal CA1 region, we observed epileptiform discharges in 6- to 7-month-old PPT1-KI mice (Fig. S5)(Musto et al., 2015). Further analysis of the neural circuitry of 1- to 2-month-old PPT1-KI mice showed that power spectral density (PSD) had an increased power of θ (3–8 Hz) and γ (30–80 Hz) bands compared to in WT mice, which could be suppressed by oral treatment with BuHA (Fig. 5A–C and E–G). Given the abnormal inhibitory neurotransmission, neural network activity, and altered expression of GABA_A_R in the PPT1-KI mice, we further explored the function of GABA_A_R as a PPT1 substrate. By genetically mutating 165 and 179 cysteines of GABA_A_R α1 to serine, we generated *Gabra1^em1^* mutant mice (Figs. S6 and S7). Remarkably, *Gabra1^em1^* mice also exhibited enhanced power in θ and γ bands (Fig. 5D, H, and I-K) and disrupted phase locking (Figs. 6D, H, I and S8D, H, I) similar to that observed in PPT1-KI mice (Fig. 5B and F). BuHA treatment partially rescued the disrupted phase locking of the theta (Fig. 6) and gamma rhythms (Fig. S8) in PPT1-KI mice. This implies that an interruption of GABA_A_R α1 subunit palmitoylation status disturbed temporal alignment.

**Figure 5.**
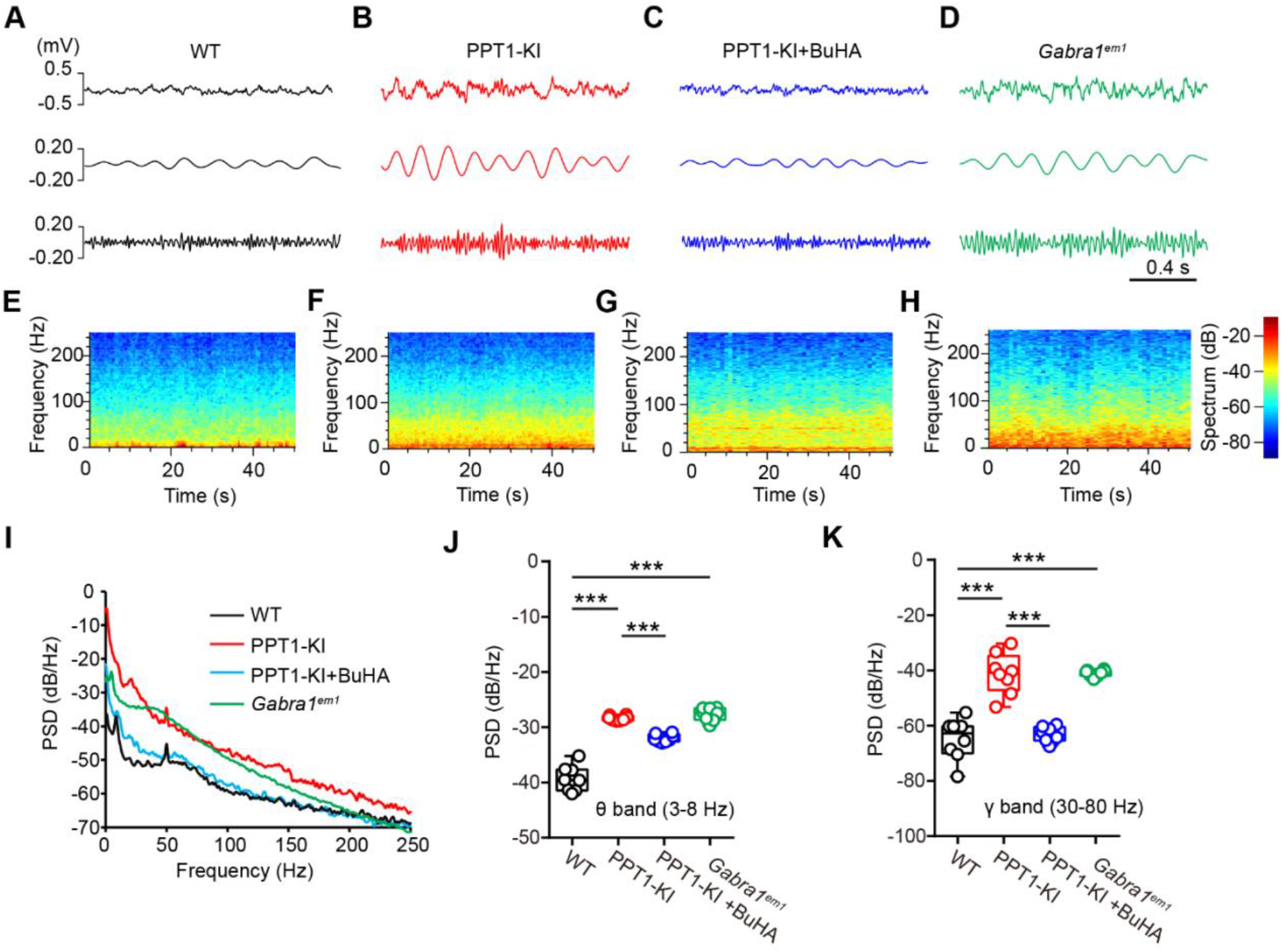
PPT1 deficiency enhances theta and gamma oscillation in CA1 region of young mice. **(A-D)** FP signals recorded in the CA1 region of WT **(A)**, PPT1-KI **(B),** PPT1-KI treated with BuHA **(C)**, and *Gabra1^em1^* mice (1 to 2 months old) **(D)**. Lower traces show filtered theta (3–8 Hz) and gamma oscillation (30–80 Hz) from the FP signals. **(E-H)** Spectrograms of the FP signals recorded from WT **(E),** PPT1-KI **(F),** BuHA-treated PPT1-KI **(G)**, and *Gabra1^em1^***(H)** mice. **(I)** PSD of FP recorded from WT (black), PPT1-KI (red), BuHA-treated PPT1-KI (blue), and *Gabra1^em1^* (green) mice. Analysis of the theta **(J)** and gamma **(K)** PSD. WT: n= 8 mice; PPT1-KI: n= 8 mice; PPT1-KI treated with BuHA: n= 8 mice; *Gabra1^em1^*: n= 6 mice; one-way ANOVA, \*\*\**P*< 0.001. Data are represented as mean ± SEM.

**Figure 6.**
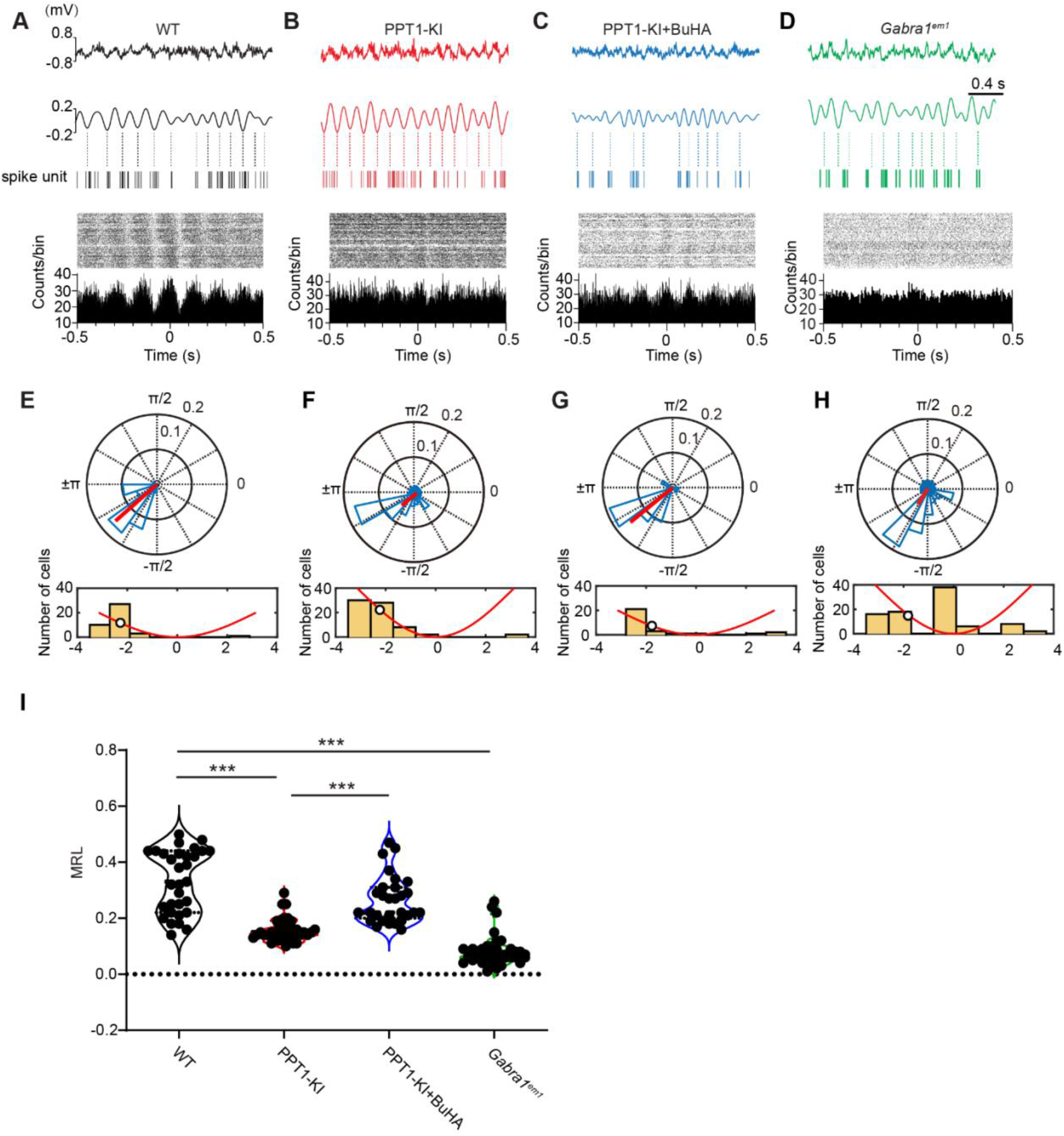
PPT1 deficiency disrupts theta phase coupling. **(A-D)** Sample traces show theta phase locking in the CA1 region. Peri-event raster (upper panel) and histogram (lower panel) displaying phase coupling of spike units and theta waves recorded from WT (**A**), PPT1-KI (**B**), BuHA-treated PPT1-KI (**C**), and *Gabra1^em1^* (**D**) mice. Tips display the timestamps of the spike units. Dotted lines indicate valleys of theta waves. Bin width is 2 ms. **(E-H)** Circular distribution of mean-spike theta phase angles (15° bin width) (upper panel) recorded from the CA1 area of WT **(E)**, PPT1-KI **(F)**, BuHA-treated PPT1-KI **(G)**, and *Gabra1^em1^* **(H)** mice. The red bars represent the direction and magnitude (length) of the MRL of the population. Distribution of mean-spike theta phase angles (45° bin width) (lower panel). The red curve displays a schematic of the theta cycle, and the white circle represents the mean phase angle. WT: −130.09°± 7.85°; PPT1-KI: −127.05°± 10.74°; PPT1- KI with BuHA: −99.99°± 17.76°; *Gabra1^em1^*: −90.72°± 11.13°; Watson–Williams test, WT vs. PPT1-KI, \**P*= 0.038; PPT1-KI vs. PPT1-KI with BuHA, *P*= 0.80; WT vs. *Gabra1^em1^*, *P*= 0.45. **(I)** Comparison of mean MRL values between multiple groups. WT: n= 31 neurons from 6 mice; PPT1- KI: n= 35 neurons from 5 mice; BuHA-treated PPT1-KI: n= 29 neurons from 5 mice; *Gabra1^em1^*: n= 40 neurons from 6 mice. Kruskal–Wallis test, \*\*\**P*< 0.001.

Research has shown that spike-phase coupling can predict memory strength(Rutishauser et al., 2010), with a recent study demonstrating that spike–phase locking is the most sensitive index for the detection of early pathological AD(Arroyo-García et al., 2021). The polar histograms indicate that most phase-locked neurons fire during the descending phase of the theta oscillations (Fig. 6E–H). The length of the mean resultant vector (MRL) is calculated to evaluate the degree of phase locking. As shown in Figure 6I, the MRL was shorter in PPT1-KI and *Gabra1^em1^* mice than in WT mice. Similarly, the gamma band polar histograms showed dramatic changes in the preferred phase and shorter MRL in PPT1-KI hippocampal CA1 neurons (Fig. S8).

These results indicate that the interruption of the GABA_A_R palmitoylation status alters theta/gamma band oscillations and uncouples the temporal relationships between neuronal discharge and theta/gamma rhythms. Our data provide pivotal evidence that normal neural activity is maintained through the PPT1-GABA_A_R α1 subunit axis, and dysfunction of such an axis may lead to neural network disorders.

### Interruption of GABA_A_R palmitoylation homeostasis impairs learning and memory in mice

Disruption of PPT1 or its binding site to the GABA_A_R α1 subunit causes massive synaptic and neural network disorders. It would be interesting to understand how learning and memory are affected in these mice. A recent study reported that PPT1-KO mice exhibit impaired long- and short-term potentiation at 3-6 months (Sapir *et al*., 2019). We found that long-term potentiation (LTP) induced in the hippocampal SC-CA1 region was normal in 1- to 2-month-old mice, but impaired in 6- to 7-month-old PPT1-KI mice likely due to the hippocampal NMDAR 2a/b and PSD95 expression decrease (Fig. S9). Consistently, the activation and expression of voltage-gated calcium channels was not altered in 1- to 2-month-old PPT1-KI mice, which sustained normal LTP formation (Fig. S10).

Despite normal LTP formation, PPT1-KI mice began to suffer from impaired spatial learning and memory as early as 2 months. As shown in Fig. 7A, PPT1-KI and *Gabra1^em1^*mice took more time to find the hidden platform and spent less time in the target entrance area in the water maze test on training days 2-5 compared to their WT littermates, and oral administration of BuHA (0.5 mM) improved their learning ability (Fig. 7A and B). No obvious differences in motor activity were detected between all the groups because swimming speed to locate the submerged escape platform during the 5 days of testing were the same (Fig. 7C). On the test day (day 6), the entry time and distance spent in the target arena were strongly reduced in PPT1-KI mice, which could be recovered by BuHA treatment (Fig. 7D–F). In addition, our Y-maze assay showed that 1- to 2-month-old PPT1-KI mice had impaired short-term spatial working memory as they entered and spent less time in the novel arm zone (Fig. 7G–I). *Gabra1^em1^* mice showed similar behavioral results compared to PPT1-KI mice (Fig. 7).

**Figure 7.**
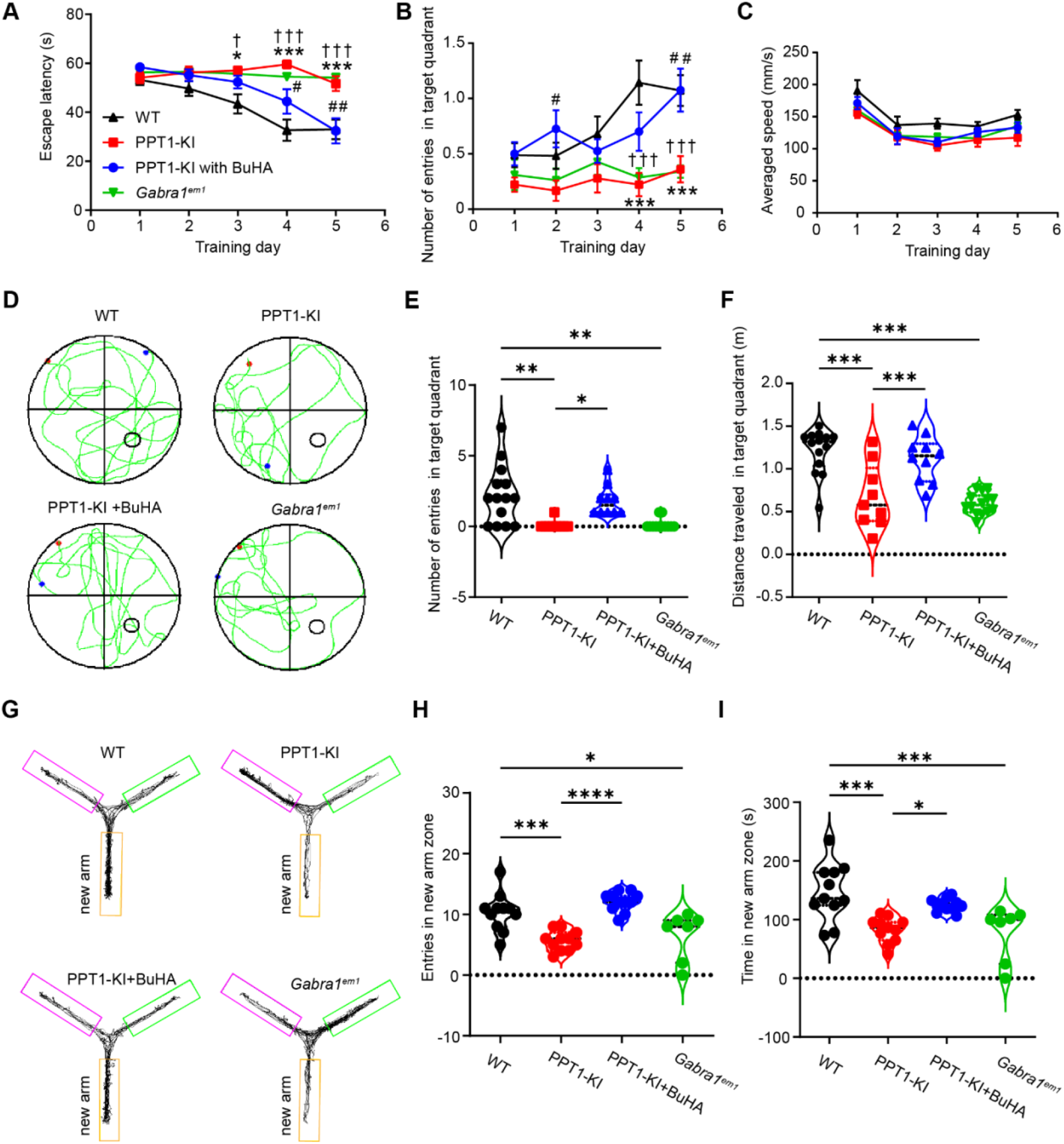
PPT1 deficiency in mice causes spatial learning and memory deficits. Average latency to reach the hidden platform **(A)**, number of times entering the platform area **(B)**, averaged swimming speed **(C)** in five days. WT (n = 14), PPT1-KI (n = 9), PPT1-KI mice treated with BuHA (n = 10), and *Gabra1^em1^* mice (n=12), two-way ANOVA, WT vs. PPT1-KI: \**P*< 0.05, \*\**P*< 0.01, \*\*\**P*< 0.001; PPT1-KI vs. PPT1-KI mice treated with BuHA: ^#^*P*< 0.05, ^##^*P*< 0.01; WT vs. *Gabra1^em1^*: ^†^*P*< 0.05, ^†††^*P*< 0.001. **(D)** Representative paths of WT, PPT1-KI, PPT1-KI mice treated with BuHA, and *Gabra1^em1^* mice were recorded on the spatial probe test day. Times entering the platform area **(E)** and total distance **(F)** in the test trial (Day 6). WT mice: n=14; PPT1-KI mice: n= 9; PPT1-KI mice treated with BuHA: n= 10; *Gabra1^em1^*mice: n=12, one-way ANOVA, \**P*< 0.05, \*\**P*< 0.01, \*\*\**P*< 0.001. **(G)** Path taken in the Y-maze test by WT, PPT1-KI, PPT1-KI mice treated with BuHA and *Gabra1^em1^*mice. Times of novel arm entries in test trial **(H)** and time spent in the novel arm zone **(I)**. WT (n = 11), PPT1-KI (n = 11), PPT1-KI mice treated with BuHA (n = 11), and *Gabra1^em1^* mice (n=7); one-way ANOVA, \**P*< 0.05, \*\*\**P*< 0.001. Data are represented as mean ± SEM.

Our results indicate that PPT1-KI and *Gabra1^em1^*mice begin to exhibit impaired spatial learning and memory formation during early development, likely due to the interruption of GABA_A_R α1 subunit palmitoylation and its resulting neural network disorders.

## Discussion

Our study identified the GABA_A_R α1 subunit as a substrate of PPT1. The palmitoylation status of the GABA_A_R α1 subunit regulates inhibitory synaptic transmission, neuronal circuit oscillation, and learning and memory ability in young mice. Our study sheds light on the potential role of postsynaptic GABA_A_Rs in the early onset of CLN1 disease.

### GABAergic neurotransmission dysfunction and neuronal apoptosis

As a fatal infantile neurodegenerative disorder, CLN1 disease causes severe nervous system deficits, such as complete retinal degeneration and blindness, cognitive and motor deficits, seizures, and flat electroencephalograms(Mole et al., 2005; Santavuori *et al*., 1973). In PPT1-deficient mice, the loss of GABAergic interneurons in the cortical region occurs before the onset of seizure activity(Kielar *et al*., 2007; Koster *et al*., 2019). Other studies have shown that GABA_A_R dysfunction is closely linked to neuroapoptosis (Hann; Qiu et al., 2015).

Similarly, we observed considerable neuron loss in the hippocampus of 6- to 7-month-old PPT1-KI mice(Zhang *et al*., 2022), as well as reduced expression of NMDAR 2a/b and PSD95, which may be the underlying mechanism for impaired LTP in old mice (Fig. S9). Neuron loss was not apparent 1-2-month-old PPT1-KI mice; however, overactivation of GABA_A_R was detected. To our surprise, newly generated *Gabra1^em1^* mice also exhibited severe developmental deficits with ∼ 80% embryonic lethality, median lifespan of 40.5 days for male, 41 days for female, and prominent weight loss irrespective of gender (Fig. S6B and C). Therefore, it is likely that the palmitoylation status change of GABA_A_R triggers chronic neuron loss in PPT1 deficiency and *Gabra1^em1^*mice, which further develops into severe symptoms, such as seizures. It would be of great interest to verify this proposal in the future.

### Palmitoylation status of GABA_A_Rα1 as a regulator of synaptic function

More than 600 palmitoylated proteins have been identified, a high percentage of which are localized to synapses(Collins et al., 2017; Kang et al., 2008; Segal-Salto et al., 2017). Approximately 10% of palmitoylated synaptic proteins are the substrates of PPT1(Gorenberg *et al*., 2022). PPT1 deficiency causes abnormal and persistent membrane accumulation of presynaptic vesicle proteins such as VAMP 2 and syntaxin-1(Kim et al., 2008); however, none of them are critical in the genesis of CLN1 disease. Proteomic studies have also predicted that many proteins involved in inhibitory GABAergic transmission, such as GAD 1/2, GABA transporter 1, and dynamin 1, are putative substrates of PPT1(Gorenberg *et al*., 2022). Numerous studies have demonstrated that both GABA_A_R(Fang et al., 2006; Keller et al., 2004; Rathenberg et al., 2004) and gephyrin(Dejanovic et al., 2014; Shen et al., 2019) are palmitoylated proteins (Fig. S3). Palmitoylation regulates the clustering and cell surface stability of GABA_A_R(Rathenberg *et al*., 2004). We discovered that PPT1 can depalmitoylate GABA_A_R α1 subunit at C260 (Fig. 4C, D). The lack of PPT1 caused abnormal membrane accumulation of GABA_A_R and enhancement of the amplitude and frequency of mIPSCs (Figs. 1 and 3D-E).

In contrast to inhibitory synapses, in PPT1-KI mice, there was no change in excitatory synaptic transmission (Fig. 2). We detected no retention of iGluRs or their scaffold proteins in the membranes of the hippocampal neurons of PPT1-KI mice at an early stage (Fig. S1). The ABE results imply that excitatory glutamate receptors are not depalmitoylated by PPT1 and are unlikely to be key molecules in the early onset of PPT1-related disease. It is also noteworthy that the mRNA expression of ABDH17B, another depalmitoylase in the central nervous system(Kanadome et al., 2019; Lin and Conibear, 2015a; Won et al., 2018; Yokoi *et al*., 2016), was lower in the hippocampus of PPT1-KI mice (Fig. S4H). Similarly, a previous study indicated that GluN2A and PSD-95 are not PPT1 substrates (Koster *et al*., 2019). Nevertheless, they demonstrated that PPT1 controls the developmental NMDAR subunit switch from GluN2B to GluN2A, probably by the depalmitoylation of Fyn kinase or GluN2B, and thus, the surface retention of GluN2B. However, it is not clear whether GluN2B is a PPT1 substrate, and why GluN2A and PSD-95 expression levels decrease from P33 to P60, whereas GluN2B remains the same (Koster *et al*., 2019). In addition, the rise time of NMDAR-mediated EPSCs did not show much difference in PPT1^-/-^ mice compared to that of the EPSC trace (Koster *et al*., 2019). Thus, PPT1 can dramatically alter synaptic strength through GABA_A_R rather than through excitatory synaptic proteins.

### Palmitoylation homeostasis of GABA_A_R regulates neural circuitry oscillation and brain function

GABA_A_R plays a critical role in generating theta and gamma oscillations, which are key to cognitive processes(Kahana et al., 1999) (Belchior et al., 2014) and are associated with cognitive related behaviours such as place coding(Buzsáki, 2005; Jensen and Lisman, 2000), memory formation(Jones and Wilson, 2005; Winson, 1978), exploratory locomotion(Buzsáki, 2002), attention, object recognition, and working memory(Buzsáki and Wang, 2012; Colgin and Moser, 2010; Garner et al., 2005; Lu et al., 2013). Further, GABA_A_R play a pivotal role in the generation of both theta and gamma oscillations(Buzsáki and Wang, 2012; Colgin and Moser, 2010; Garner *et al*., 2005; Lu and Henderson, 2010; Lu *et al*., 2013; Soltesz and Deschênes, 1993). Our *in vivo* FP recordings indicated enhanced theta and gamma oscillations, but loss of theta or gamma phase locking in the CA1 area of 1- to 2-month-old PPT1-deficit mice under free-moving conditions. However, the substrates and downstream signalling pathways that mediate abnormalities in brain oscillations remain unclear. Among the hundreds of neuronal targets of PPT1, we found that GABA_A_R is one of the substrates. Interestingly, when we generated *Gabra1^em1^* mice by mutating the binding sites between PPT1 and GABA_A_R α1 at C165 and C179, *in vivo* FP data also showed enhanced theta and gamma oscillations in the CA1 area, as well as poor phase-locking in theta or gamma waves, which is similar to *CLN1*^-/-^(PPT1-KI) mice (Figs. 5, 6, and Fig. S8).

This provides the missing evidence that the GABA_A_R α1 subunit is the targeting molecules in PPT1-induced neurological disorders. Moreover, PPT1 deficiency induces serious epilepsy in both humans suffering from NCLs(Kravljanac and Sims, 2021; Maeser et al., 2021; Schulz et al., 2013) and animals(Gupta et al., 2001). In PPT1^-/-^ mice, seizures can be detected after 7 months using electroencephalogram recordings(Kielar *et al*., 2007). Consistently, we observed two types of epileptiform discharges at 6–7 months old but not in 1- to 2-month-old PPT1-KI mice (Fig. S5). It is possible that the potentiation of low-frequency oscillations with the chaos of neuronal discharges recorded in early stage gradually developed into epileptiform discharges at later stages.

Neural circuitry oscillations are fundamental to higher brain functions. One study found that older *CLN1*^−/−^ mice (5–6 months) exhibited impairments in spatial learning and memory (Dearborn et al., 2015). Similarly, our data showed that learning and memory were impaired, even in 1- to 2-month-old PPT1-KI and *Gabra1^em1^* mice (Fig. 7), which strongly correlates with the disrupted oscillation waves. Surprisingly, these memory deficits in PPT1-KI mice could be partially rescued by feeding the mice BuHA, indicating the crucial role of PPT1 in disease genesis.

In summary, we found that the GABA_A_R α1 subunit is a depalmitoylation substrate of PPT1, and its palmitoylation homeostasis determines GABAergic neurotransmission, neuronal network oscillations, and high brain function in mice. Mutation of GABA_A_R and PPT1 binding sites mimicked the phenotype of PPT1-KI mice, highlighting the importance of GABA_A_R in the pathogenesis of NCL. These findings present GABA_A_R as a therapeutic target that can be leveraged not only for CLN1 disease but also for other GABA_A_R dysfunction-induced neurodegenerative disorders.

## Acknowledgments

This work was supported by funds from the Scientific and Technological Project in He’nan Province 232102311227 (JT), Open Project Program of the Third Affiliated Hospital of Xinxiang Medical University KFKTYB202111 (JT), Startup Grants of Xinxiang Medical University 300-505481 (LHG), National Natural Science Foundation 81771517 (CBL), Hubei 3551 Talented Fund 3551-T82 (SYP). We thank Hai Huang and Wenyan Han for helpful comments on this manuscript.

## Author contributions

Conceived studies: SP, LG, JT, and CL

Generated GABAAR α1 mutant mice: SP and LG

Conducted review and editing: LG, JT, SP, CL and JH

Performed the experiments or data analysis: JT, JG, YQ, YL, ZG, XZ, QW, GY, ZL, JL, HS, JL, YM, YL, JX and MR

Provided funding acquisition, project administration and resources: LG, JT, SP, and CL

Wrote the paper: LG, JT and SP

All the authors have read and approved the final version of the manuscript.

## Data availability statement

All data are available in the main text or the supplementary materials.

## Study approval

Animal experiments were performed according to the regulations and requirements of the XXMU Animal Ethics Committee (No. XYLL2021053).

## Materials and Methods

### Animals

#### _C57BL/6N-_^Gabra1em1Cin(p.C165S,p.C179S)^_/Cya (Gabra1_^em1^_) mutant mice_

The mouse *Gabra1* gene (GenBank accession number: NM_010250.5; Ensembl: ENSMUSG00000010803) is located on chromosome 11. Ten exons were identified, with an ATG start codon in exon 2 and a TAG stop codon in exon 10. p.C165 and p.C179 are located in exon 6. Exon 6 was selected as the target site. We designed a gRNA targeting vector and donor oligo (with a targeting sequence flanked by 240 bp homologous sequences combined on both sides) (**Table 1**).

**Table 1.**
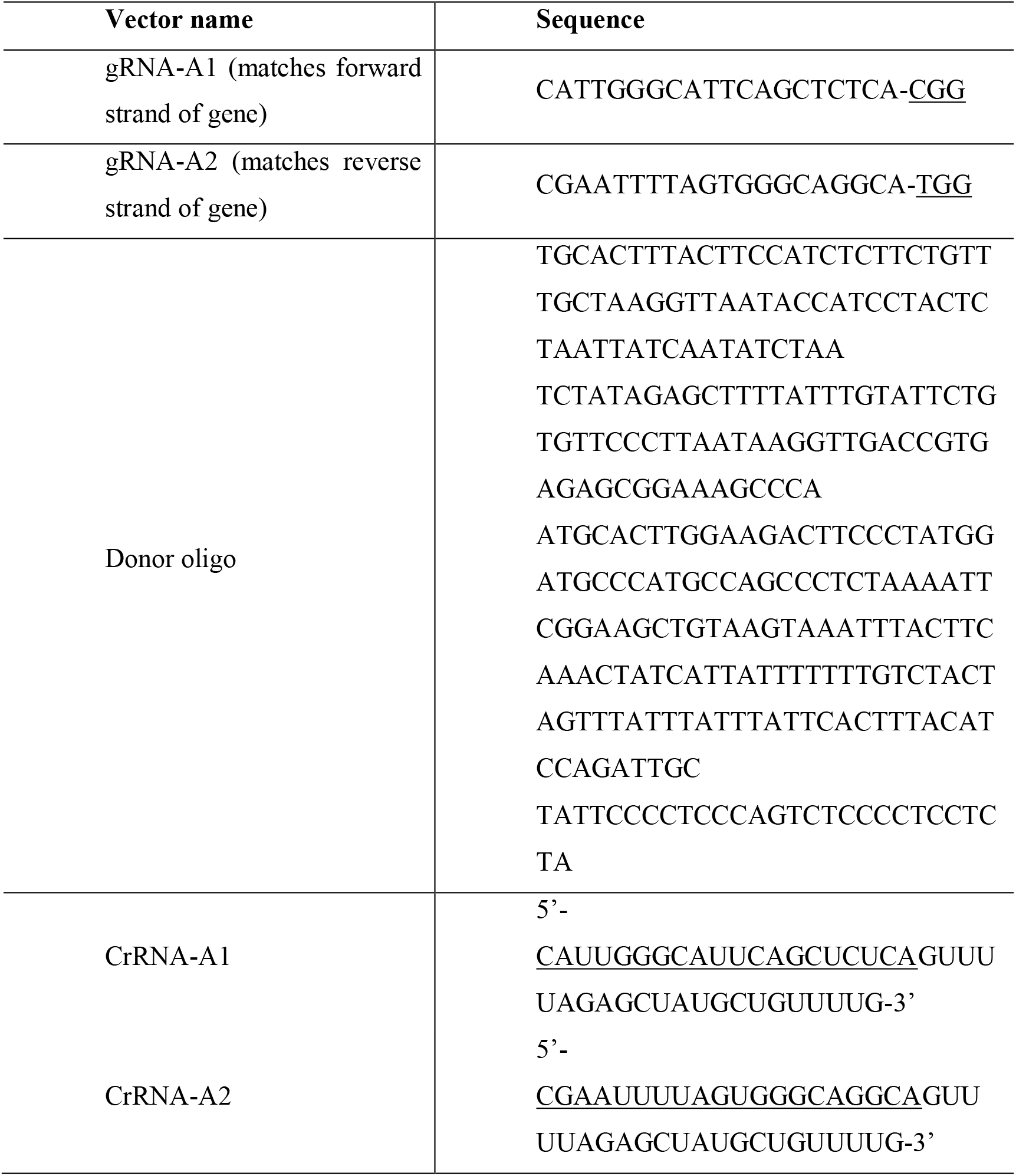
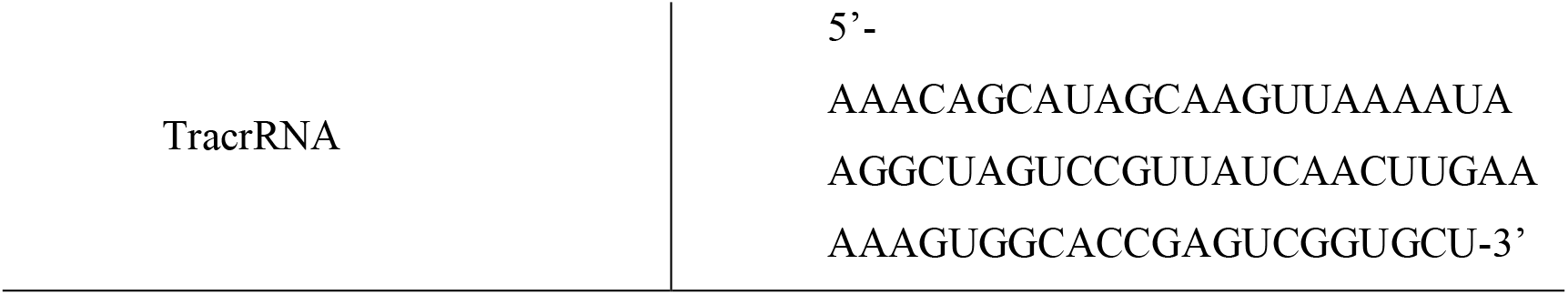
Nuclease expression vector for *Gabra1* point mutation knock-in.

The gRNA to the mouse *Gabra1* gene, the donor oligo containing p.C165S (TGC to AGC) and p.C179S (TGC to AGC) mutations, two synonymous mutations (p.A163= (GCT to GCG) and p.P180= (CCA to CCT)), and Cas9 were co-injected into fertilized mouse eggs, followed by implantation of the eggs into surrogate mothers to obtain offspring to generate targeted knock-in offspring. F0 founder animals were identified by PCR, followed by sequence analysis, and were bred with WT mice to test germline transmission and F1 animal generation. F2 homozygotes were generated by F1 autocopuation. The target region of the mouse *Gabra1* locus was amplified by PCR using specific primers. The PCR products were sequenced to confirm the targeting. Primer sequence: Primer-F: 5’-GGATTAGTGGGTGCTCTCTGTTAT-3’; Primer-R: 5’-GCCTGTGGATCTTCTACCCTAAC-3’.

### PPT1-KI (CLN1 c.451C>T (p.R151X)) mutant mice

The strategy for point mutation of the *CLN1* gene to generate PPT-KI mice has been described as before. As previous studies, PPT1-KI mice carrying c.451C>T/c.451C>T mutations are deficient in PPT1-protein as well as PPT1 enzyme activity. And the PPT1-KI mice did not show structural deterioration of the retinal layers until 6 months of age(Bouchelion *et al*., 2014). C57BL/6N mice were purchased from Beijing Vital River Laboratory Animal Technology Co., Ltd, Beijing, China (animal licence number:2016-0006) and used as the WT controls. All animals were housed and maintained in the specific pathogen-free animal facility of the Animal Experiment Center of the Institute of Psychiatry and Neuroscience of Xinxiang Medical University (XXMU) under a 12-h light/dark cycle. Animals had ad libitum access to food and water, except during food and water deprivation periods. A previous study demonstrated that BuHA can easily cross the blood-brain barrier in the mouse brain(Sarkar *et al*., 2013). PPT1-KI mice and their littermates were orally administered 1 mM BuHA for 1 month before mating to breast-feed pups with BuHA after parturition. Young pups had free access to food and water (containing 1 mM BuHA) throughout the experiments. All efforts were made to minimise animal suffering and reduce the number of animals used. Experiments were performed in accordance with the guidelines published in the NIH Guide for the Care and Use of Laboratory Animals (NIH publication no. 86-23, revised 1987). All reagents were purchased from Sigma-Aldrich (St. Louis, MO, USA), unless otherwise stated.

#### Electrophysiology

### In vivo electrophysiological recording

#### Stereotaxic surgery and electrode implantation

Mice were anesthetised with an intraperitoneal injection of 1% pentobarbital sodium (0.45– 0.5 mL/100 g). Under anaesthesia, mice were secured in a stereotaxic frame with ear bars. The head was shaved with a razor, and a midline 5-mm incision was made using a sterile scalpel. The subcutaneous tissue was removed from the skull, and a craniotomy (∼1.5 × 0.5 mm) was drilled (AP: 1.82 mm, ML:1.25 mm, DV:1.5 mm, right hemisphere) in the CA1 region. Two steel screws are anchored to the anterior and posterior edges of the surgical site to secure the implant. After the endocranium was removed, a 4 × 2 micro wire electrode (KD-MWA-8, 25 μm nitinol wire, Kedou (Suzhou) Brain-Computer Technology Co., Ltd.) with 3 μm polyethylene glycol coating was implanted into the pyramidal cell layer of CA1. The craniotomy site was sealed with a sterile silicone elastomer (Kwik-Sil WPI) to prevent brain injury. After surgery, the implanted electrodes and screws were cemented integrally to the skull using a denture base resin type II (Shanghai Medical Instruments Co., Ltd.). After surgery, the animals were housed individually on a reversed 12/12 h day/night schedule.

#### Signal acquisition

Following one week of recovery, wideband signals were recorded using an OmniPlex Neural Recording Data Acquisition System (Plexon Inc., Dallas, TX, USA) with an 8 kHz global low-pass filter. A continuous spike was sampled at 4 kHz, followed by a 300 Hz low-cut filter. The FP was set to a 200 low-pass filter and down-sampled to 1 kHz. After recording, the hippocampus was post-fixed for Nissl staining to verify the proper placement of the electrodes in the target region.

#### Spike sorting

Spikes were sorted using Offline Sorter (Plexon Inc., Dallas, TX, USA) to classify the electrical activity of individual neurons, based on the first to third principal components(Adamos et al., 2008). Spike units were excluded when the absolute refractory period of single-unit autocorrelation was < 1 ms. Cross-channel artefacts identified by their time coincidence across channels were also invalidated.

#### Spectral analysis

The power spectrum density and spectrogram of continuous FP were computed using NeuroExplorer (Nex Technologies, Colorado Springs, CO, USA) with 1024 frequency values and 25% window overlap. Before this process, the FP signal values were multiplied by the coefficients of the Hann window and discrete fast Fourier transformations of the results were calculated using formulas defined previously(Sfondouris et al., 2012). Theta/gamma waves were filtered by bandpass filtering of the FP data using NeuroExplorer software with Digital Filtering of Continuous Variables function.

#### Phase locking analysis

To evaluate the wave phase locking, waves were further processed through Hilbert transformation using built-in or custom-built MATLAB scripts (R2016a, MathWorks, USA). Only neurons with ≥ 50 spikes during the period analysed were used for phase locking estimation. We tested the significance of spike-FP phase locking using circular statistics (CirStat toolbox in Matlab)(Berens, 2009; Fisher and N., 1993; Gregoire, 1996). Rayleigh’s test was employed to assess the circular distribution of the mean phase angle and to test the nonuniformity of each neuron’s spike phase distribution to theta or gamma. Neurons were considered significantly phase-locked at *P*< 0.05. The mean phase angle was computed in the circular direction of the mean resultant vector. The Watson-Williams *F*-test was performed to compare the mean phase angles of neurons recorded from multiple groups.

We evaluated the phase locking of neurons by calculating the mean resultant vector (MRL, range 0–1) length of the spike phase angle distribution. An MRL value of one indicates exact phase synchrony, whereas a value of zero indicates no phase synchrony. The non-parametric Kruskal– Wallis test was performed to test the differences in the MRL values between multiple groups. *P*< 0.05 was considered statistically significant. Moreover, the valley of the wave timestamp was identified as a reference event using Find Oscillation function to plot the peri-event raster. Data and MATLAB scripts supporting the findings of this study are available from the corresponding author upon request.

### In vitro electrophysical recording

#### Slice preparation

The mice were anaesthetised with urethane and perfused with ice-cold artificial cerebrospinal fluid (aCSF) through the left ventricle until the limbs turned white. The brain was then rapidly removed and immersed in ice-cold aCSF-containing 225 mM sucrose, 3 mM KCl, 6 mM MgCl_2_, 1.25 mM NaH_2_PO_4_, 24 mM NaHCO_3_, 0.5 mM CaCl_2_, and 10 mM glucose. Transverse slices (350 μM thickness) were prepared using a vibratome (Ci-7000SMZ2, Campden instrument, Loughborough, UK). Immediately after preparation, slices were transferred to a nylon net within a chamber, and two sides of the chamber were exposed to normal aCSF containing mM 126 NaCl, 3 mM KCl, 1.25 mM NaH_2_PO_4_, 2 mM MgSO_4_, 24 mM NaHCO_3_, 2 mM CaCl_2_, and 10 mM glucose, bubbled with a mixture of 95 % O_2_ and 5 % CO_2_, and kept at pH 7.35-7.45 at room temperature (RT) for storage or 32 °C for recording. For the BuHA treatment group, slices were incubated with 1 μM BuHA for 1 h at 37 ℃ before patch clamp recording. The perfusion rate was maintained at 1-2 mL/min (Guo et al., 2017).

#### LTP induction

We stimulated the Schaffer collateral (SC) projecting from CA3 to the pyramidal layer of CA1 and extracellularly recorded the field excitatory postsynaptic potentials (fEPSPs). To determine the standard stimulation intensity for each slice, stimuli were administered at varying intensities, and the PSP amplitude was used to construct a stimulus-response curve. For SC-CA1 LTP recording, the fEPSPs were evoked in the stratum radiatum layer at CA1 using glass pipettes (3–4 MΩ) by stimulating the SC of CA3 with a concentric bipolar microelectrode (25 μm inner pole diameter, 30203, FHC, Bowdoin, ME). Pulses of 0.1 ms duration were delivered every 30 s. The standard stimulus intensity was set 40% of the maximum fEPSP slope. An fEPSP amplitude of less than 1 mV before tetanus indicated an unhealthy condition of slice and was excluded from the experiment. LTP was induced by a double high-frequency stimulation (100 Hz for 1 s). Recordings were performed at 32 °C.

#### Patch clamp recording

We measured mIPSCs in voltage-clamp mode at −70 mV in the presence of a metabotropic glutamate receptors blocker, [250 μM (S)-α-methyl-4-carboxyphenylglycine (MCPG)], GABA_B_ receptor blocker (1 μM CGP55845), and 1 μM tetrodotoxin (TTX)(O’Neill and Sylantyev, 2018). The pipette solution contained 140 mM CsCl, 10 mM Na-HEPES, 10 mM EGTA, 2 mM Mg-ATP, and 5 mM QX-314 (pH 7.3). To block EPSCs, 25 μM D-(–)-2-amino-5-phosphonopentanoic acid (D-APV) and 5 μM 6-cyano-7-nitroquinoxaline-2,3-dione (CNQX) were added in the bath solution just before use(Chakrabarti et al., 2016; Sabanov et al., 2017). For the evoked IPSCs recordings, the cells were clamped at +40 mV(Kapur et al., 1997). All patch-clamp recordings were performed at RT.

To isolate AMPAR-mediated miniature excitatory postsynaptic currents (mEPSCs), 1 μM TTX, 10 μM D-APV, and 10 μM bicuculline were added to the bath solution. Pipette (3-4 MΩ) solution contained 125 mM Cs-gluconate, 20 mM KCl, 4 mM Mg-ATP, 10 mM Na_2_-phosphocreatine, 0.3 mM Na_2_-GTP, 10 mM HEPES, 0.5 mM EGTA, and 5 mM QX314 (pH 7.3) adjusted with CsOH. For evoked EPSC recording, neurons were voltage-clamped at −70 mV to record AMPAR-mediated EPSCs or at +40 mV to record dual-component EPSCs containing NMDAR-mediated EPSCs. AMPAR/NMDAR ratios were also calculated by dividing the peak of the AMPAR-mediated EPSC at −70 mV by the value of the NMDAR-mediated EPSC after a stimulation start time of 50 ms at +40 mV.

For the isolation of voltage-dependent inward calcium currents, 1 μM TTX and 20 mM TEA-Cl were added to the bath solution to block Na^+^ and K^+^ channels. The recipe of the pipette solution was similar to that of the postsynaptic pipette solution, except that K^+^ was replaced with Cs^+^. Cells were clamped at −80 mV and then depolarised to +60 mV for 20 ms in 10 mV steps.

Currents were amplified using a Multiclamp 700 B amplifier (Molecular Devices, Sunnyvale, CA, USA), low-pass filtered at 1 kHz, and digitised using a Digidata 1550B interface (Molecular Devices) at 5 kHz. Synaptic currents were detected and analysed using pClamp 10 software (Molecular Devices). Offline analysis of mEPSCs/IPSCs was performed using Clampfit 10.4.2.0 (Molecular Devices) software. Only recordings where series resistance remained below 16 MΩ and did not increase by more than 20 % during the experiment were included in the analysis (Sabanov *et al*., 2017).

### Cell culture and Immunostaining

Primary cultures of hippocampal neurons were prepared as previously described (Thomas et al., 2012). Briefly, timed-pregnant mice at E17.5–18.5 were anaesthetised on ice and decapitated. The foetal hippocampi were quickly dissected out in an ice-cold Hank’s balanced salt solution without calcium or magnesium, containing 1 mM HEPES (GIBCO; ThermoFisher Scientific, Waltham, MA) and penicillin (100 units/mL) or streptomycin (100 μg/mL) (GIBCO). The hippocampi were triturated with a sterile tweezer and digested in Hanks’ balanced salt solution containing 0.25% trypsin and 0.1 mg/mL DNase Ⅰ at 37 ℃ for 15 min. The digested hippocampus was minced in DMEM with 5% FBS culture medium and filtered through a 70 μm cell strainer. The filtrate was centrifuged at 500 rpm for 10 min and resuspended in neurobasal feeding medium containing 2% B27 supplement and 2 mM glutamine solution. The cells were plated onto poly-D-lysine (100 μg/mL, Sigma) and laminin (4 mg/mL, Sigma)-coated glass coverslips and incubated at 37 ℃ with 5% CO_2_.

Cells were washed three times with PBS and fixed with 4% paraformaldehyde or methanol at DIV15. Then cells were permeabilized with 0.1% Triton X-100 in PBS. After blocking with 3% BSA, the cells were incubated with primary antibodies overnight at 4 ℃ followed by Alexa Fluor-conjugated secondary antibodies (Abcam) for 1 h at RT. The primary antibodies used were anti-PPT1 (DF7733; Affinity) and anti-GABAA1 (ab94585; Abcam). The secondary antibodies were goat anti-rabbit IgG H&L (Alexa Fluor®594) and goat anti-mouse IgG H&L (Alexa Fluor® 647). After DAPI counterstaining, cells were mounted using Fluoromount-G™ Slide Mounting Medium (EMS, 17984-25), and fluorescence was visualised by confocal microscopy (Leica TCS SP8 STED, Germany) at a dark place.

HEK293T cell lines were obtained from ATCC (Manassas, VA). The cells were cultured in high-glucose Dulbecco’s Modified Eagle Media (DMEM) (Servicebio, cat: G4515-500 mL) with 10 % (v/v) fetal bovine serum (FBS) (Invigentech, Cat: A6903 FBS-500) and 1 % (v/v) penicillin-streptomycin solution (Hyclone, SV30010) at 37 °C with 5 % CO_2_.

### STED image

The cells were imaged using 594 and 647 nm excitation lasers coupled with STED 775 nm depletion lasers. XY/Z_stack_ scan mode with the logical size of 1024×1024 (15 nm×15 nm pixels per voxel size) and scan speed at 600 Hz was set to capture the entire cell images under 100 x oil objective, numerical aperture: 1.4 with 151.6 μm pinhole. All images were deconvolved using Huygens software (Scientific Volume Imaging, Hilversum, Holland) and analysed using the LAS X software (Leica). Colocalisation was analysed using Image J 6.0 software (National Institutes of Health, Bethesda, MD, USA) with the colocalization analysis plugin (Bolte and Cordelières, 2006).

### ABE assay

We performed the ABE as previously described (Hayashi et al., 2009; Wan et al., 2007). Briefly, the lysates were incubated with 10 mM N-ethylmaleimide (NEM, E3876; Sigma) overnight at 4 °C. NEM was then removed using three sequential chloroform/methanol (CM) precipitation steps. After three CM precipitations, proteins were solubilised in solubilising buffer containing 1 M hydroxylamine hydrochloride (159417, Sigma), 1 mM HPDP-biotin (A8008; APExBIO, Houston, TX), 0.2% TritonX-100 (T8787, Sigma), and protease inhibitors (4693132001; Roche, Basel, Switzerland) in phosphate-buffered saline (pH 7.4). After incubation for 1 h at RT, proteins were precipitated by CM and then suspended with 200 μM HPDP-biotin and 0.2% Triton X-100 for 1.5–2 h at RT. After 3 CM precipitations, the proteins were incubated with streptavidin-agarose (16-126; Millipore sigma, Burlington, MA) for one night at 4 °C. The beads were washed five times with wash buffer containing 150 mM NaCl, 50 mM Tris-HCl, 5 mM EDTA (pH 7.4), and 0.2% Triton X-100. After washing, the proteins were eluted with wash buffer containing 1.5% β-mercaptoethanol (Sigma-Aldrich, M3148) at 37 °C for 0.5 h with agitation (350 rpm) and then heated in a 100 °C water bath for 10 min. After centrifugation, the released proteins in the supernatant were denatured in sodium dodecyl sulphate sample buffer and processed for western blotting with primary antibody ABE assays. All other biochemical experiments were performed at least three times. In each case, a representative experiment is presented.

The potential palmitoylation sites (C165, C179, C260, C319) of the GABA_A_R α1 subunit were forecast using the CSS-Palm Online Service webset. All mutants (GABA_A_Rα1 C165A, C179A, C165A&179A, C260A&C319A) with Myc tags were purchased from Fenghui Biology (Hu’nan, China). The ABE assay was performed as described previously.

Other experimental procedures are provided in the **Supplemental Information:**

***S1 Behavioural studies***

Morris water maze (MWM) Y maze

***S2 Biochemical analysis***

Cell membrane/cytoplasmic protein extraction Co-immunoprecipitation (Co-IP)

Western blot

Quantitative polymerase chain reaction (qPCR) Cell culture

### Statistics

All data were acquired and analysed by researchers who were blinded to the genotype of the mice and acute slices. All data were analysed using SPSS Statistics 20 (IBM, Armonk, NY, USA). We confirmed the homogeneity of variances using Levene’s test and the equality of means using the Brown–Forsythe test. Data were statistically analysed by Student’s *t*-test for two-group comparisons and one-way analysis of variance (ANOVA) for multi-group comparisons. Data that did not follow a normal distribution were estimated by the non-parametric Mann–Whitney *U*-test for two-group comparisons and the Kruskal–Wallis test for multi-group comparisons. Cumulative probability distribution analysis was performed using the Kolmogorov-Smirnov test. Differences in the mean phase angles of multiple groups of neurons were evaluated using Watson-Williams *F*-test. Western blot results were analysed using the t-test for two-group comparisons and one-way ANOVA for multi-group comparisons.

## References

Adamos, D.A., Kosmidis, E.K., and Theophilidis, G. (2008). Performance evaluation of PCA-based spike sorting algorithms. Computer methods and programs in biomedicine 91, 232–244. 10.1016/j.cmpb.2008.04.011.

Ahrens-Nicklas, R.C., Tecedor, L., Hall, A.F., Lysenko, E., Cohen, A.S., Davidson, B.L., and Marsh, E.D. (2019). Neuronal network dysfunction precedes storage and neurodegeneration in a lysosomal storage disorder. JCI insight 4. 10.1172/jci.insight.131961.

Arroyo-García, L.E., Isla, A.G., Andrade-Talavera, Y., Balleza-Tapia, H., Loera-Valencia, R., Alvarez-Jimenez, L., Pizzirusso, G., Tambaro, S., Nilsson, P., and Fisahn, A. (2021). Impaired spike-gamma coupling of area CA3 fast-spiking interneurons as the earliest functional impairment in the App(NL-G-F) mouse model of Alzheimer’s disease. Mol Psychiatry 26, 5557–5567. 10.1038/s41380-021-01257-0.

Belchior, H., Lopes-Dos-Santos, V., Tort, A.B., and Ribeiro, S. (2014). Increase in hippocampal theta oscillations during spatial decision making. Hippocampus 24, 693–702. 10.1002/hipo.22260.

Berens, P.J.J.o.S.S. (2009). CircStat: A MATLAB Toolbox for Circular Statistics. 31.

Bolte, S., and Cordelières, F.P. (2006). A guided tour into subcellular colocalization analysis in light microscopy. J Microsc 224, 213–232. 10.1111/j.1365-2818.2006.01706.x.

Bouchelion, A., Zhang, Z., Li, Y., Qian, H., and Mukherjee, A.B. (2014). Mice homozygous for c.451C>T mutation in Cln1 gene recapitulate INCL phenotype. Ann Clin Transl Neurol 1, 1006–1023. 10.1002/acn3.144.

Buzsáki, G. (2002). Theta oscillations in the hippocampus. Neuron 33, 325–340. 10.1016/s0896-6273(02)00586-x.

Buzsáki, G. (2005). Theta rhythm of navigation: link between path integration and landmark navigation, episodic and semantic memory. Hippocampus 15, 827–840. 10.1002/hipo.20113.

Buzsáki, G., and Wang, X.J. (2012). Mechanisms of gamma oscillations. Annual review of neuroscience 35, 203–225. 10.1146/annurev-neuro-062111-150444.

Camp, L.A., Verkruyse, L.A., Afendis, S.J., Slaughter, C.A., and Hofmann, S.L. (1994). Molecular cloning and expression of palmitoyl-protein thioesterase. The Journal of biological chemistry 269, 23212–23219.

Chakrabarti, S., Qian, M., Krishnan, K., Covey, D.F., Mennerick, S., and Akk, G. (2016). Comparison of Steroid Modulation of Spontaneous Inhibitory Postsynaptic Currents in Cultured Hippocampal Neurons and Steady-State Single-Channel Currents from Heterologously Expressed alpha1beta2gamma2L GABA(A) Receptors. Molecular pharmacology 89, 399–406. 10.1124/mol.115.102202.

Chu-LaGraff, Q., Blanchette, C., O’Hern, P., and Denefrio, C. (2010). The Batten disease Palmitoyl Protein Thioesterase 1 gene regulates neural specification and axon connectivity during Drosophila embryonic development. PloS one 5, e14402. 10.1371/journal.pone.0014402.

Colgin, L.L., and Moser, E.I. (2010). Gamma oscillations in the hippocampus. Physiology (Bethesda, Md.) 25, 319–329. 10.1152/physiol.00021.2010.

Collins, M.O., Woodley, K.T., and Choudhary, J.S. (2017). Global, site-specific analysis of neuronal protein S-acylation. Scientific reports 7, 4683. 10.1038/s41598-017-04580-1.

Dearborn, J.T., Harmon, S.K., Fowler, S.C., O’Malley, K.L., Taylor, G.T., Sands, M.S., and Wozniak, D.F. (2015). Comprehensive functional characterization of murine infantile Batten disease including Parkinson-like behavior and dopaminergic markers. Scientific reports 5, 12752. 10.1038/srep12752.

Dejanovic, B., Semtner, M., Ebert, S., Lamkemeyer, T., Neuser, F., Lüscher, B., Meier, J.C., and Schwarz, G. (2014). Palmitoylation of gephyrin controls receptor clustering and plasticity of GABAergic synapses. PLoS biology 12, e1001908. 10.1371/journal.pbio.1001908.

Fang, C., Deng, L., Keller, C.A., Fukata, M., Fukata, Y., Chen, G., and Lüscher, B. (2006). GODZ-mediated palmitoylation of GABA(A) receptors is required for normal assembly and function of GABAergic inhibitory synapses. The Journal of neuroscience : the official journal of the Society for Neuroscience 26, 12758–12768. 10.1523/jneurosci.4214-06.2006.

Fisher, and N., I. (1993). Statistical Analysis of Circular Data || Descriptive methods. 10.1017/CBO9780511564345, 15–38.

Garner, H.L., Whittington, M.A., and Henderson, Z. (2005). Induction by kainate of theta frequency rhythmic activity in the rat medial septum-diagonal band complex in vitro. The Journal of physiology 564, 83–102. 10.1113/jphysiol.2004.080622.

Gorenberg, E.L., Massaro Tieze, S., Yücel, B., Zhao, H.R., Chou, V., Wirak, G.S., Tomita, S., Lam, T.T., and Chandra, S.S. (2022). Identification of substrates of palmitoyl protein thioesterase 1 highlights roles of depalmitoylation in disulfide bond formation and synaptic function. PLoS biology 20, e3001590. 10.1371/journal.pbio.3001590.

Gregoire, T.G.J.F.S. (1996). Statistical Analysis of Circular Data. 4.

Guo, F., Zhao, J., Zhao, D., Wang, J., Wang, X., Feng, Z., Vreugdenhil, M., and Lu, C. (2017). Dopamine D4 receptor activation restores CA1 LTP in hippocampal slices from aged mice. Aging Cell 16, 1323–1333. 10.1111/acel.12666.

Gupta, P., Soyombo, A.A., Atashband, A., Wisniewski, K.E., Shelton, J.M., Richardson, J.A., Hammer, R.E., and Hofmann, S.L. (2001). Disruption of PPT1 or PPT2 causes neuronal ceroid lipofuscinosis in knockout mice. Proc Natl Acad Sci U S A 98, 13566–13571. 10.1073/pnas.251485198.

Hann, U.A.V.a.S.R. The a1 Subunit of GABAA Receptor Is Repressed by c-Myc and is Pro-Apoptotic. Journal of Cellular Biochemistry.

Hayashi, T., Thomas, G.M., and Huganir, R.L. (2009). Dual palmitoylation of NR2 subunits regulates NMDA receptor trafficking. Neuron 64, 213–226. 10.1016/j.neuron.2009.08.017.

Hobert, J.A., and Dawson, G. (2006). Neuronal ceroid lipofuscinoses therapeutic strategies: past, present and future. Biochimica et biophysica acta 1762, 945–953. 10.1016/j.bbadis.2006.08.004.

Jensen, O., and Lisman, J.E. (2000). Position reconstruction from an ensemble of hippocampal place cells: contribution of theta phase coding. Journal of neurophysiology 83, 2602–2609. 10.1152/jn.2000.83.5.2602.

Jin, J., Zhi, X., Wang, X., and Meng, D. (2021). Protein palmitoylation and its pathophysiological relevance. Journal of cellular physiology 236, 3220–3233. 10.1002/jcp.30122.

Jones, M.W., and Wilson, M.A. (2005). Theta rhythms coordinate hippocampal-prefrontal interactions in a spatial memory task. PLoS biology 3, e402. 10.1371/journal.pbio.0030402.

Kahana, M.J., Sekuler, R., Caplan, J.B., Kirschen, M., and Madsen, J.R. (1999). Human theta oscillations exhibit task dependence during virtual maze navigation. Nature 399, 781–784. 10.1038/21645.

Kanadome, T., Yokoi, N., Fukata, Y., and Fukata, M. (2019). Systematic Screening of Depalmitoylating Enzymes and Evaluation of Their Activities by the Acyl-PEGyl Exchange Gel-Shift (APEGS) Assay. Methods in molecular biology (Clifton, N.J.) 2009, 83–98. 10.1007/978-1-4939-9532-5_7.

Kang, R., Wan, J., Arstikaitis, P., Takahashi, H., Huang, K., Bailey, A.O., Thompson, J.X., Roth, A.F., Drisdel, R.C., Mastro, R., et al. (2008). Neural palmitoyl-proteomics reveals dynamic synaptic palmitoylation. Nature 456, 904–909. 10.1038/nature07605.

Kapur, A., Pearce, R.A., Lytton, W.W., and Haberly, L.B. (1997). GABAA-mediated IPSCs in piriform cortex have fast and slow components with different properties and locations on pyramidal cells. Journal of neurophysiology 78, 2531–2545. 10.1152/jn.1997.78.5.2531.

Keller, C.A., Yuan, X., Panzanelli, P., Martin, M.L., Alldred, M., Sassoè-Pognetto, M., and Lüscher, B. (2004). The gamma2 subunit of GABA(A) receptors is a substrate for palmitoylation by GODZ. The Journal of neuroscience : the official journal of the Society for Neuroscience 24, 5881–5891. 10.1523/jneurosci.1037-04.2004.

Kielar, C., Maddox, L., Bible, E., Pontikis, C.C., Macauley, S.L., Griffey, M.A., Wong, M., Sands, M.S., and Cooper, J.D. (2007). Successive neuron loss in the thalamus and cortex in a mouse model of infantile neuronal ceroid lipofuscinosis. Neurobiol Dis 25, 150–162. 10.1016/j.nbd.2006.09.001.

Kim, S.J., Zhang, Z., Sarkar, C., Tsai, P.C., Lee, Y.C., Dye, L., and Mukherjee, A.B. (2008). Palmitoyl protein thioesterase-1 deficiency impairs synaptic vesicle recycling at nerve terminals, contributing to neuropathology in humans and mice. The Journal of clinical investigation 118, 3075–3086. 10.1172/jci33482.

Koster, K.P., Francesconi, W., Berton, F., Alahmadi, S., Srinivas, R., and Yoshii, A. (2019). Developmental NMDA receptor dysregulation in the infantile neuronal ceroid lipofuscinosis mouse model. eLife 8. 10.7554/eLife.40316.

Kravljanac, R., and Sims, K. (2021). A case of juvenile CLN1-challenge in diagnosis and epilepsy treatment. Neurocase 27, 165–168. 10.1080/13554794.2021.1905852.

Lange, J., Haslett, L.J., Lloyd-Evans, E., Pocock, J.M., Sands, M.S., Williams, B.P., and Cooper, J.D. (2018). Compromised astrocyte function and survival negatively impact neurons in infantile neuronal ceroid lipofuscinosis. Acta neuropathologica communications 6, 74. 10.1186/s40478-018-0575-4.

Lemonidis, K., Werno, M.W., Greaves, J., Diez-Ardanuy, C., Sanchez-Perez, M.C., Salaun, C., Thomson, D.M., and Chamberlain, L.H. (2015). The zDHHC family of S-acyltransferases. Biochemical Society transactions 43, 217–221. 10.1042/bst20140270.

Lin, D.T., and Conibear, E. (2015a). ABHD17 proteins are novel protein depalmitoylases that regulate N-Ras palmitate turnover and subcellular localization. eLife 4, e11306. 10.7554/eLife.11306.

Lin, D.T., and Conibear, E. (2015b). Enzymatic protein depalmitoylation by acyl protein thioesterases. Biochemical Society transactions 43, 193–198. 10.1042/bst20140235.

Lu, C.B., and Henderson, Z. (2010). Nicotine induction of theta frequency oscillations in rodent hippocampus in vitro. Neuroscience 166, 84–93. 10.1016/j.neuroscience.2009.11.072.

Lu, C.B., Li, C.Z., Li, D.L., and Henderson, Z. (2013). Nicotine induction of theta frequency oscillations in rodent medial septal diagonal band in vitro. Acta pharmacologica Sinica 34, 819–829. 10.1038/aps.2012.198.

Maeser, S., Petre, B.A., Ion, L., Rawer, S., Kohlschütter, A., Santorelli, F.M., Simonati, A., Schulz, A., and Przybylski, M. (2021). Enzymatic diagnosis of neuronal lipofuscinoses in dried blood spots using substrates for concomitant tandem mass spectrometry and fluorimetry. Journal of mass spectrometry : JMS 56, e4675. 10.1002/jms.4675.

Mitchell, D.A., Vasudevan, A., Linder, M.E., and Deschenes, R.J. (2006). Protein palmitoylation by a family of DHHC protein S-acyltransferases. Journal of lipid research 47, 1118–1127. 10.1194/jlr.R600007-JLR200.

Mole, S.E., Williams, R.E., and Goebel, H.H. (2005). Correlations between genotype, ultrastructural morphology and clinical phenotype in the neuronal ceroid lipofuscinoses. Neurogenetics 6, 107–126. 10.1007/s10048-005-0218-3.

Mukherjee, A.B., Appu, A.P., Sadhukhan, T., Casey, S., Mondal, A., Zhang, Z., and Bagh, M.B. (2019). Emerging new roles of the lysosome and neuronal ceroid lipofuscinoses. Molecular neurodegeneration 14, 4. 10.1186/s13024-018-0300-6.

Musto, A.E., Walker, C.P., Petasis, N.A., and Bazan, N.G. (2015). Hippocampal neuro-networks and dendritic spine perturbations in epileptogenesis are attenuated by neuroprotectin d1. PloS one 10, e0116543. 10.1371/journal.pone.0116543.

Nakazono, T., Jun, H., Blurton-Jones, M., Green, K.N., and Igarashi, K.M. (2018). Gamma oscillations in the entorhinal-hippocampal circuit underlying memory and dementia. Neuroscience research 129, 40–46. 10.1016/j.neures.2018.02.002.

O’Neill, N., and Sylantyev, S. (2018). Spontaneously opening GABA(A) receptors play a significant role in neuronal signal filtering and integration. Cell death & disease 9, 813. 10.1038/s41419-018-0856-7.

Qiu, J., Shi, P., Mao, W., Zhao, Y., Liu, W., and Wang, Y. (2015). Effect of apoptosis in neural stem cells treated with sevoflurane. BMC Anesthesiol 15, 25. 10.1186/s12871-015-0018-8.

Rathenberg, J., Kittler, J.T., and Moss, S.J. (2004). Palmitoylation regulates the clustering and cell surface stability of GABAA receptors. Molecular and cellular neurosciences 26, 251–257. 10.1016/j.mcn.2004.01.012.

Rutishauser, U., Ross, I.B., Mamelak, A.N., and Schuman, E.M. (2010). Human memory strength is predicted by theta-frequency phase-locking of single neurons. Nature 464, 903–907. 10.1038/nature08860.

Sabanov, V., Braat, S., D’Andrea, L., Willemsen, R., Zeidler, S., Rooms, L., Bagni, C., Kooy, R.F., and Balschun, D. (2017). Impaired GABAergic inhibition in the hippocampus of Fmr1 knockout mice. Neuropharmacology 116, 71–81. 10.1016/j.neuropharm.2016.12.010.

Santavuori, P., Haltia, M., Rapola, J., and Raitta, C. (1973). Infantile type of so-called neuronal ceroid-lipofuscinosis. 1. A clinical study of 15 patients. Journal of the neurological sciences 18, 257–267. 10.1016/0022-510x(73)90075-0.

Santorelli, F.M., Garavaglia, B., Cardona, F., Nardocci, N., Bernardina, B.D., Sartori, S., Suppiej, A., Bertini, E., Claps, D., Battini, R., et al. (2013). Molecular epidemiology of childhood neuronal ceroid-lipofuscinosis in Italy. Orphanet journal of rare diseases 8, 19. 10.1186/1750-1172-8-19.

Sapir, T., Segal, M., Grigoryan, G., Hansson, K.M., James, P., Segal, M., and Reiner, O. (2019). The Interactome of Palmitoyl-Protein Thioesterase 1 (PPT1) Affects Neuronal Morphology and Function. Frontiers in cellular neuroscience 13, 92. 10.3389/fncel.2019.00092.

Sarkar, C., Chandra, G., Peng, S., Zhang, Z., Liu, A., and Mukherjee, A.B. (2013). Neuroprotection and lifespan extension in Ppt1(-/-) mice by NtBuHA: therapeutic implications for INCL. Nature neuroscience 16, 1608–1617. 10.1038/nn.3526.

Schulz, A., Kohlschütter, A., Mink, J., Simonati, A., and Williams, R. (2013). NCL diseases - clinical perspectives. Biochimica et biophysica acta 1832, 1801–1806. 10.1016/j.bbadis.2013.04.008.

Segal-Salto, M., Hansson, K., Sapir, T., Kaplan, A., Levy, T., Schweizer, M., Frotscher, M., James, P., and Reiner, O. (2017). Proteomics insights into infantile neuronal ceroid lipofuscinosis (CLN1) point to the involvement of cilia pathology in the disease. Human molecular genetics 26, 1678. 10.1093/hmg/ddx074.

Sfondouris, J.L., Quebedeaux, T.M., Holdgraf, C., and Musto, A.E. (2012). Combined process automation for large-scale EEG analysis. Computers in biology and medicine 42, 129–134. 10.1016/j.compbiomed.2011.10.017.

Shen, Z.C., Wu, P.F., Wang, F., Xia, Z.X., Deng, Q., Nie, T.L., Zhang, S.Q., Zheng, H.L., Liu, W.H., Lu, J.J., et al. (2019). Gephyrin Palmitoylation in Basolateral Amygdala Mediates the Anxiolytic Action of Benzodiazepine. Biological psychiatry 85, 202–213. 10.1016/j.biopsych.2018.09.024.

Sleat, D.E., Gedvilaite, E., Zhang, Y., Lobel, P., and Xing, J. (2016). Analysis of large-scale whole exome sequencing data to determine the prevalence of genetically-distinct forms of neuronal ceroid lipofuscinosis. Gene 593, 284–291. 10.1016/j.gene.2016.08.031.

Soltesz, I., and Deschênes, M. (1993). Low- and high-frequency membrane potential oscillations during theta activity in CA1 and CA3 pyramidal neurons of the rat hippocampus under ketamine-xylazine anesthesia. Journal of neurophysiology 70, 97–116. 10.1152/jn.1993.70.1.97.

Sung-Jo Kim, Louis Dye, Anil B. Mukherjee Palmitoyl protein thioesterase-1 deficiency impairs synaptic vesicle recycling at nerve terminals, contributing to neuropathology in humans and mice.

Thomas, G.M., Hayashi, T., Chiu, S.L., Chen, C.M., and Huganir, R.L. (2012). Palmitoylation by DHHC5/8 targets GRIP1 to dendritic endosomes to regulate AMPA-R trafficking. Neuron 73, 482–496. 10.1016/j.neuron.2011.11.021.

Vesa, J., Hellsten, E., Verkruyse, L.A., Camp, L.A., Rapola, J., Santavuori, P., Hofmann, S.L., and Peltonen, L. (1995). Mutations in the palmitoyl protein thioesterase gene causing infantile neuronal ceroid lipofuscinosis. Nature 376, 584–587. 10.1038/376584a0.

Wan, J., Roth, A.F., Bailey, A.O., and Davis, N.G. (2007). Palmitoylated proteins: purification and identification. Nature protocols 2, 1573–1584. 10.1038/nprot.2007.225.

Williams, R.E. (2011). 361Appendix 1: NCL Incidence and Prevalence Data. In The Neuronal Ceroid Lipofuscinoses (Batten Disease), S. Mole, R. Williams, and H. Goebel, eds. (Oxford University Press), pp. 0. 10.1093/med/9780199590018.003.0023.

Winson, J. (1978). Loss of hippocampal theta rhythm results in spatial memory deficit in the rat. Science 201, 160–163. 10.1126/science.663646.

Won, S.J., Cheung See Kit, M., and Martin, B.R. (2018). Protein depalmitoylases. Critical reviews in biochemistry and molecular biology 53, 83–98. 10.1080/10409238.2017.1409191.

Yokoi, N., Fukata, Y., Sekiya, A., Murakami, T., Kobayashi, K., and Fukata, M. (2016). Identification of PSD-95 Depalmitoylating Enzymes. The Journal of neuroscience : the official journal of the Society for Neuroscience 36, 6431–6444. 10.1523/jneurosci.0419-16.2016.

Zhang, X., Wang, M., Feng, B., Zhang, Q., Tong, J., Wang, M., Lu, C., and Peng, S. (2022). Seizures in PPT1 Knock-In Mice Are Associated with Inflammatory Activation of Microglia. International journal of molecular sciences 23. 10.3390/ijms23105586.

