## supplementary information for "GABA_A_R-PPT1 palmitoylation homeostasis controls synaptic transmission and circuitry oscillation in CLN1 disease"

#### Supplementary Text

#### S1 Behavioral studies

##### Morris water maze (MWM)

PPT1-KI mice and their WT littermates were used to perform the MWM test based on previously described studies(Morris, 1984; Vorhees and Williams, 2006). Mice were trained in a circular black maze (diameter, 120 cm; depth, 50 cm) (NDS-MWM-1, Noldus, Wageningen, the Netherlands) and filled with opaque water (24 °C) made with milk. Four spaced labels around the edge of the pool were used as starting positions: east (E), southeast (SE), northwest (NW), and north (N). A platform of 10-cm diameter was placed in the center of the SW quadrant and submerged 1.2 cm under the water surface. For training period (days 1-5), each mouse underwent four trials with a maximum of 60 s/day to find the hidden platform placed at the central of NE quadrant. If the mouse located the platform before 60 s had passed, it was immediately removed from the pool. If the platform was not located after 60 s of swimming, the mouse was gently guided to the platform and allowed to re-orient to the distal visual cues for an additional 20 s before being removed from the pool. After removal from the pool, the mice were manually dried with a towel and placed in a warming cage for at least 5 min before returning to the home cage. Twenty-four hours after the last training trial (day 6), retention was assessed during the probe trial in which the platform was removed. Animals were video-tracked using Etho vision software (Noldus, Wageningen, the Netherlands), and swim speed, traveled distance, latency, and percentage of time spent in each quadrant were automatically calculated.

##### Y maze

Y maze used to assess spatial recognition memory was identical to that described by Pietropaolo et al.(Pietropaolo et al., 2009). The animals were allowed to acclimatize to the environment for 7 days. Testing was carried out using a Y-shaped maze with three light-colored, opaque arms orientated at 120° angles from each other. Mouse was placed into the Y-maze with one arm (novel arm) of the maze closed off during training session. On testing session, 30 min after the mouse was removed from the maze, the mouse was placed back into the maze with the blockage removed. The number of entries into the novel arm is then compared to the entries into the other arms to assess the degree of spatial memory. A mouse that shows no preference for any of the arms during the testing session is an indication of an impaired spatial memory, which may indicate impaired hippocampal function. Allocation of arms (start, familiar, and novel) was counterbalanced within each experimental group. Animals were video-tracked using Etho vision software (Noldus, Wageningen, the Netherlands).

**S2 Biochemical analysis**

**Cell membrane/cytoplasmic protein extraction**

The extraction procedure was performed according to the manufacturer's instructions (P0033; Beyotime Biotechnology, Shanghai, China). The primary antibodies used in this study were anti-GRIN-2b (21920, Proteintech), anti-NMDA-2a (ab124913, Abcam), anti-glutamate receptor 1 (AMPA subtype) (ab109450, Abcam), anti-PSD 95 (#2507; Cell Signaling Technology), anti-SAP102 (S19-2) (MA5-27707, Invitrogen), anti-pan cadherin [CH-19] (ab6528, Abcam), anti-PSD 93 (MABN497; Millipore Sigma), anti-GABAAR alpha1 (DF8548, Affinity Biosciences), GABAAR beta 2/GABRB2 (ab186875, Abcam), anti-GABAAR gamma 2 (DF6583, Affinity Biosciences), anti-gephyrin (12681-1-AP, Proteintech), anti-c-Myc antibody (C3956, Sigma), anti-FLAG® M2 (#8146, Cell Signaling Technology), and anti-β-actin (AC006, ABclonal). The second antibodies were HRP goat anti-mouse IgG (H+L) (AS003, ABclonal) and HRP goat anti-rabbit IgG (H+L) (AS014, ABclonal). The results were statistically analyzed using Student's t-test.

**Co-immunoprecipitation (Co-IP)**

For brain tissue lysate immunoprecipitation, hippocampi were dissected and homogenized in ice-cold IP lysis buffer. After centrifugation at 800 g for 5 min, the supernatant was collected and centrifuged at 12,000 g for 15 min. The pellet was then resuspended in the pre-chilled lysis buffer and incubated on ice for 1 hr. The hippocampal lysates were centrifuged at 12,000 g for 15 min again and the supernatant was incubated with anti-PPT1 or GABAAR a1 antibody at 4 ℃ overnight. On the next day, 50 µL of Pierce Protein A/G Magnetic Beads (Thermo, 88802) was added for 1 h at RT. The beads were collected with a magnetic stand and then washed three times with IP lysis buffer (20 mM Tris (pH 7.5), 150 mM NaCl, 1% Triton X-100, sodium pyrophosphate, β-glycerophosphate, EDTA, Na3VO4, leupeptin). The precipitated proteins were eluted with SDS loading buffer with β-mercaptoethanol and denatured at 100 ºC for 10 min before subjecting to SDS-PAGE and immunoblotting.

For cultured cells immunoprecipitation, HEK293T cells were co-transfected with PPT1-Flag and WT or mutants (C165A, C179A, C165A&179A, C260A&C319A) $GABA_AR$ α1-Myc using

Lipofectamine 2000 for 24-48 hr before harvesting with IP buffer containing protease inhibitor mixture (Roche). The insoluble debris was removed by centrifugation at 15,000 g for 10 min at 4 °C. The soluble supernatant was then separated into input and IP lysates. An equal amount of the IP lysate was incubated with 40 μL Anti-FLAG® M2 Affinity gel (Sigma-Aldrich) at 4 ºC overnight while rotating. The beads were collected by centrifugating at 800 g 5 min, and then washed three times with IP lysis buffer. As mentioned above, the precipitated proteins were eluted with SDS loading buffer with β-mercaptoethanol and denatured at 100 ºC for 10 min before subjecting to SDS-PAGE and immunoblotting.

**Western blot**

Protein was resolved by SDS-PAGE and then transferred onto PVDF membranes, blocked and incubated with primary antibodies at 4 ºC overnight. The membranes were then washed 3 times with 0.1 TBST and incubated with HRP-conjugated secondary antibodies for 1 h at RT. Protein was detected with the standard enhanced chemiluminescence (ECL) method and documented by a gel imaging system (Tanon). The following primary antibodies were used in the assay: anti-GRIN-2b (21920, Proteintech), anti-NMDA-2a (ab124913, Abcam), anti-glutamate receptor 1 (AMPA subtype) (ab109450, Abcam), anti-PSD 95 (#2507; Cell Signaling Technology), anti-SAP102 (S19-2) (MA5-27707, Invitrogen), anti-pan cadherin [CH-19] (ab6528, Abcam), anti-PSD 93 (MABN497; Millipore Sigma), anti-GABAAR alpha1 (DF8548, Affinity Biosciences), GABAAR beta 2/GABRB2 (ab186875, Abcam), anti-GABAAR gamma 2 (DF6583, Affinity Biosciences), anti-gephyrin (12681-1-AP, Proteintech), anti-c-Myc antibody (C3956, Sigma),anti-Myc (#2276; Cell Signaling Technology) , anti-FLAG® M2 (#8146, Cell Signaling Technology), and anti-β-actin (AC006, ABclonal), HRP-conjugated mouse anti-DDDDK-Tag (AE024, ABclonal) . The second antibodies were HRP goat anti-mouse IgG (H+L) (AS003, ABclonal) and HRP goat anti-rabbit IgG (H+L) (AS014, ABclonal).

**Quantitative polymerase chain reaction (qPCR)**

A FastPure Cell/Tissue Total RNA Isolation Kit (RC101-01, Vazyme) was used to extract total RNAs from the mice hippocampi, according to manufacturer's protocols. HiScript III RT SuperMix for qPCR (+gDNA wiper) (R323-01, Vazyme) was used to synthetise complementary DNA (cDNA). Relative quantitation of target gene expression was measured by the $2^{-\triangle\triangle}$Ct method and β-actin was deemed as a reference. The primer sequences used are as follows (5′ to 3′):

GABAAα1, TTCACAAGAATTTTGGACCGAC (forward) and CCTTCCAACTTTGACGGAAAAA (reverse); GABAAβ2, TACTCAGCACGCTTGAGATAAA (forward) and CAATGGCATTCACATCAGTCAA (reverse); GABAAγ2, ACACCATGCGTCTGACAATCTCTG (forward) and GCCATCTTCTGCCACCACCAC (reverse).

**Cell culture**

HEK293T cell lines were obtained from ATCC (Manassas, VA). The cells were cultured in high-glucose Dulbecco's Modified Eagle Media (DMEM) (Servicebio, cat: G4515-500 mL) with 10 % (v/v) fetal bovine serum (FBS) (Invigentech, Cat: A6903 FBS-500) and 1 % (v/v) penicillin- streptomycin solution (Hyclone, SV30010) at 37 °C with 5 % $CO_2$.

#### Supplemental figure legends

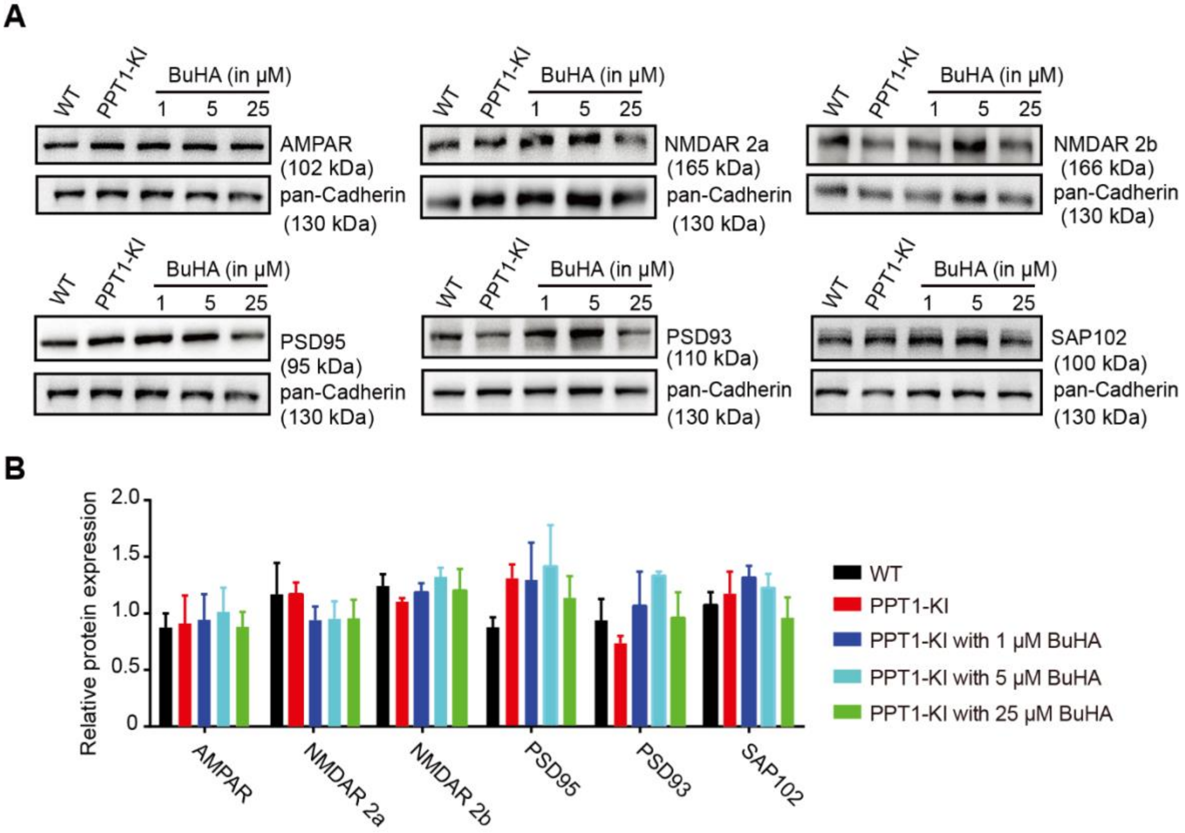


**Figure S1. Membrane expression of iGluRs and scaffold proteins are not influenced at 1- to 2- months old PPT1-KI mice.**

**(A, B)** Western blot analysis of iGluRs (AMPAR, NMDAR 2a/2b) and scaffold proteins (PSD 95/93, SAP 102) extracted from the hippocampal membrane fraction. N=3 for each group, two-way ANOVA, no significant difference. Data are represented as mean ± SEM.

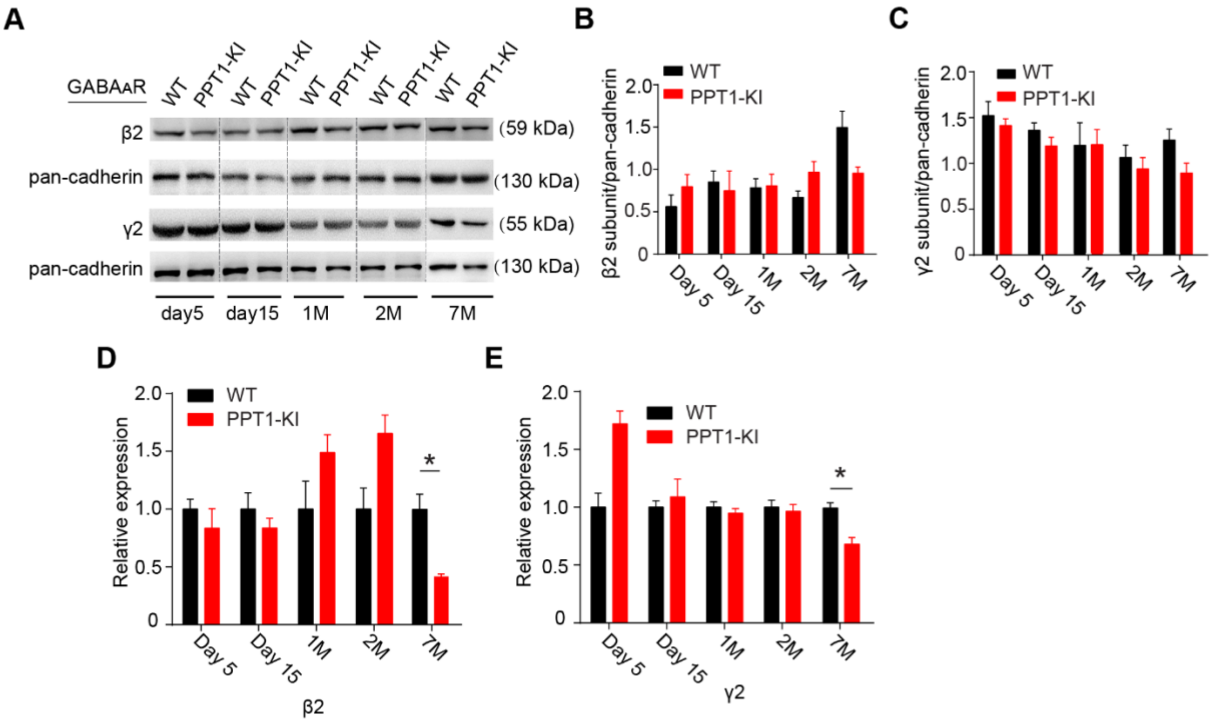


**Figure S2. Membrane expression of GABA$_A$R β2 and γ2 subunits were not developmentally regulated in PPT1-KI mice.**

**(A-D)** Representative immunoblot **(A)** and quantification of GABA$_A$R β2 **(B)** and γ2 **(C)** subunits levels on the cellular membrane in WT and PPT1-KI mice at indicated ages. N=3 for each group, two-way ANOVA, *$P$< 0.05.

**(D, E)** Quantitative PCR showing mRNA expression of GABA$_A$R β2 **(D)** and γ2 **(E)** subunits at indicated ages. N=3, two-way ANOVA, *$P$< 0.05. Data are represented as mean ± SEM.

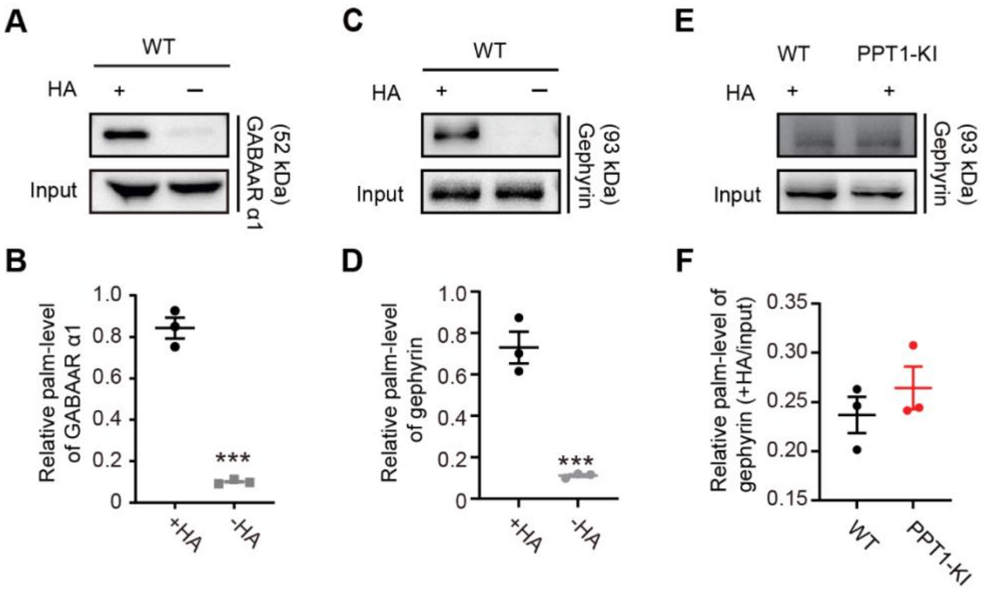


**Figure S3. PPT1 deficiency does not change gephyrin palmitoylation level.**

**(A-D)** Representative immunoblot and quantitative analysis of palmitoylated $GABA_A R$ α1 subunit **(A** and **B)** and gephyrin **(C** and **D)** measured using ABE assay in WT mice hippocampus.

**(E, F)** PPT1 deficiency did not alter palmitoylation level of gephyrin.

+HA, with hydroxylamine; n= 3 for each group, t-test, ****P*< 0.001. Data are represented as mean ± SEM.

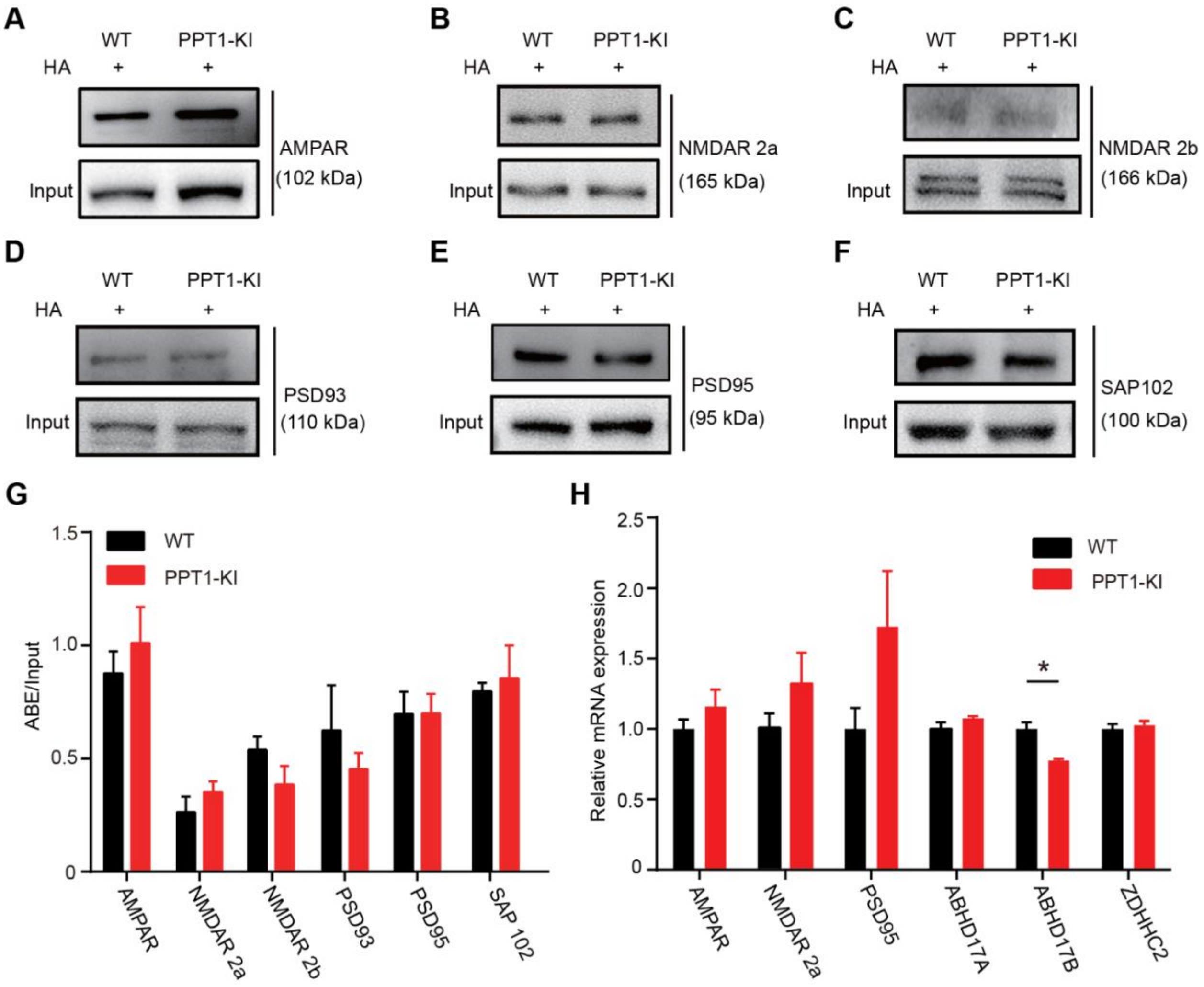


**Figure S4. Palmitoylation status of iGluRs and the scaffold proteins in the hippocampi of PPT1-KI mice.**

**(A-F)** Representative immunoblots of palmitoylated AMPAR **(A)**, NMDAR 2a **(B)**, NMDAR 2b **(C)**, and postsynaptic scaffold proteins PSD 93 **(D)**, PSD 95 **(E)**, SAP 102 **(F)**. GAPDH: glyceraldehyde-3-phosphate dehydrogenase; +HA: with hydroxylamine.

**(G)** Bar graph showing the normalised ratio of acyl biotin exchange to input. N=3 for each group, two-way ANOVA, no significant difference.

**(H)** Reverse transcription PCR results for NMDAR 2a, AMPAR, PSD 95, ABHD 17A/B, and ZDHHC2. N=3 for each group, two-way ANOVA, *$P<0.05$.

Data are represented as mean ± SEM.

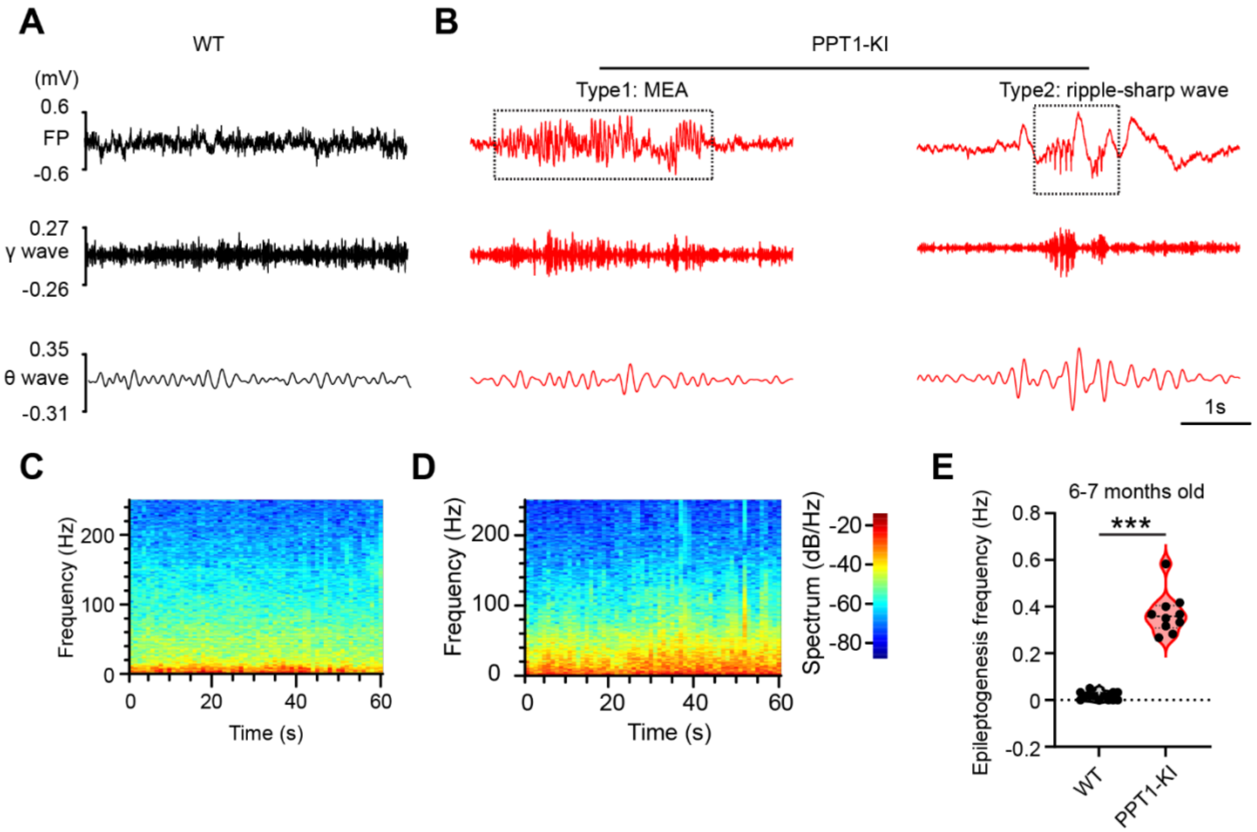


**Figure S5. Epileptiform activities in the CA1 region of 6 to 7 months old PPT1-KI mice.**

**(A, B)** LFP signals recorded at CA1 region from WT **(A)** and PPT1-KI **(B)** mice (6-7 months). The lower traces showing filtered theta (3-8 Hz) and gamma oscillation (30-80 Hz) from LFP signals. The dashed line box in **B** indicated epileptiform discharges detected from CA1 region from PPT1-KI mice. MEA: microepileptiform activity.

**(C, D)** Spectrograms of LFP signals recorded from WT **(C)** and PPT1-KI **(D)** mice.

**(E)** Analysis of epileptogenesis frequency. WT: n= 13 mice; PPT1-KI: n=10 mice, t-test, ***$P$< 0.001.

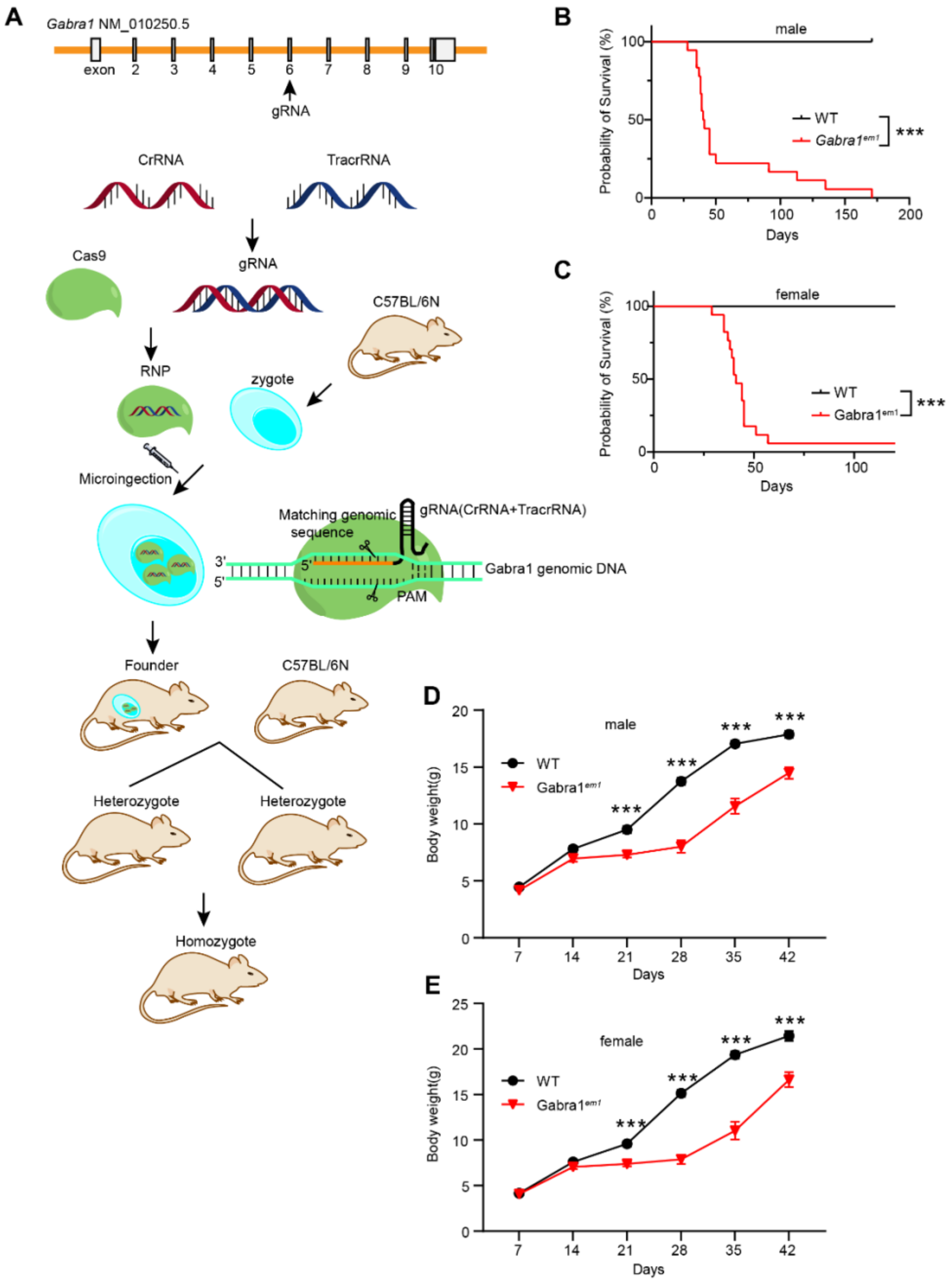


**Figure S6.** ***Gabra1*** **gene and homozygote reproduction, life-span, and body weight assess.** **(A)** Point mutation strategy design of *Gabra1* gene and homozygote reproduction. **(B, C)** Kaplan–Meier survival curve of WT (male: n = 19; female: n = 14) and *Gabra1*$^{em1}$(male: n = 35; female: n = 22). Chi-square test, ***$P$< 0.001. **(D, E)** Weight growth curve of WT (male: n = 31; female: n = 33) and *Gabra1*$^{em1}$ mice (male: n = 42; female: n = 38) (day 7 to day 42 after birth). Two-way ANOVA, ***$P$< 0.001. gRNA: guide RNA ; CrRNA: CRISPR-derived RNA; TracrRNA: trans-activating crRNA.

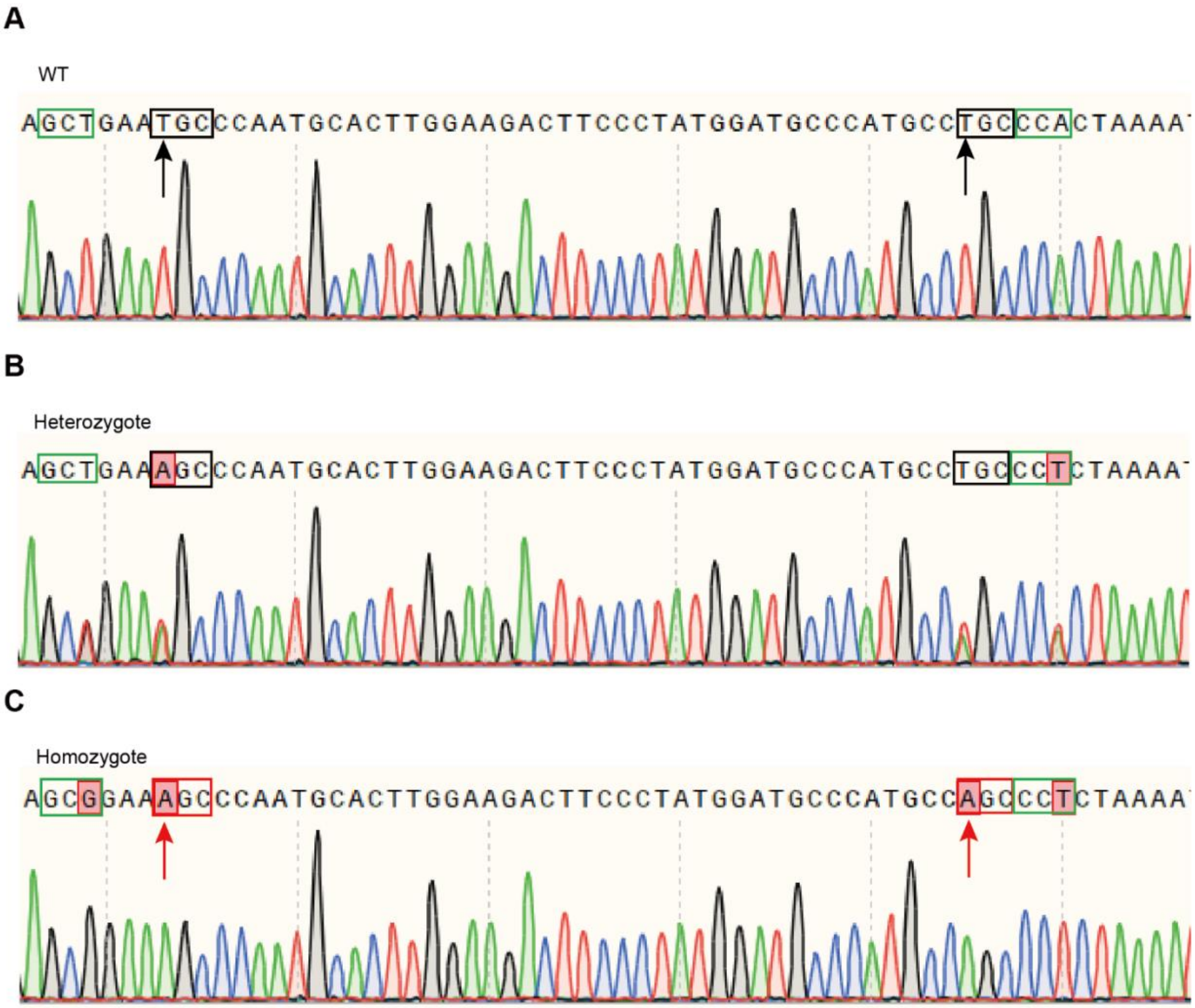


**Figure S7. Gene sequencing results of animals with p.C165S (TGC to AGC) and p.C179S (TGC to AGC) mutations and silent mutations p.A163= (GCT to GCG) and p.P180= (CCA to CCT).**

**(A)** Gene sequencing result of WT *Gabra1*.

**(B, C)** Gene sequencing result of *Gabra1* with TGC to AGC heterozygous mutation **(B)** and homozygous mutation **(C)**.

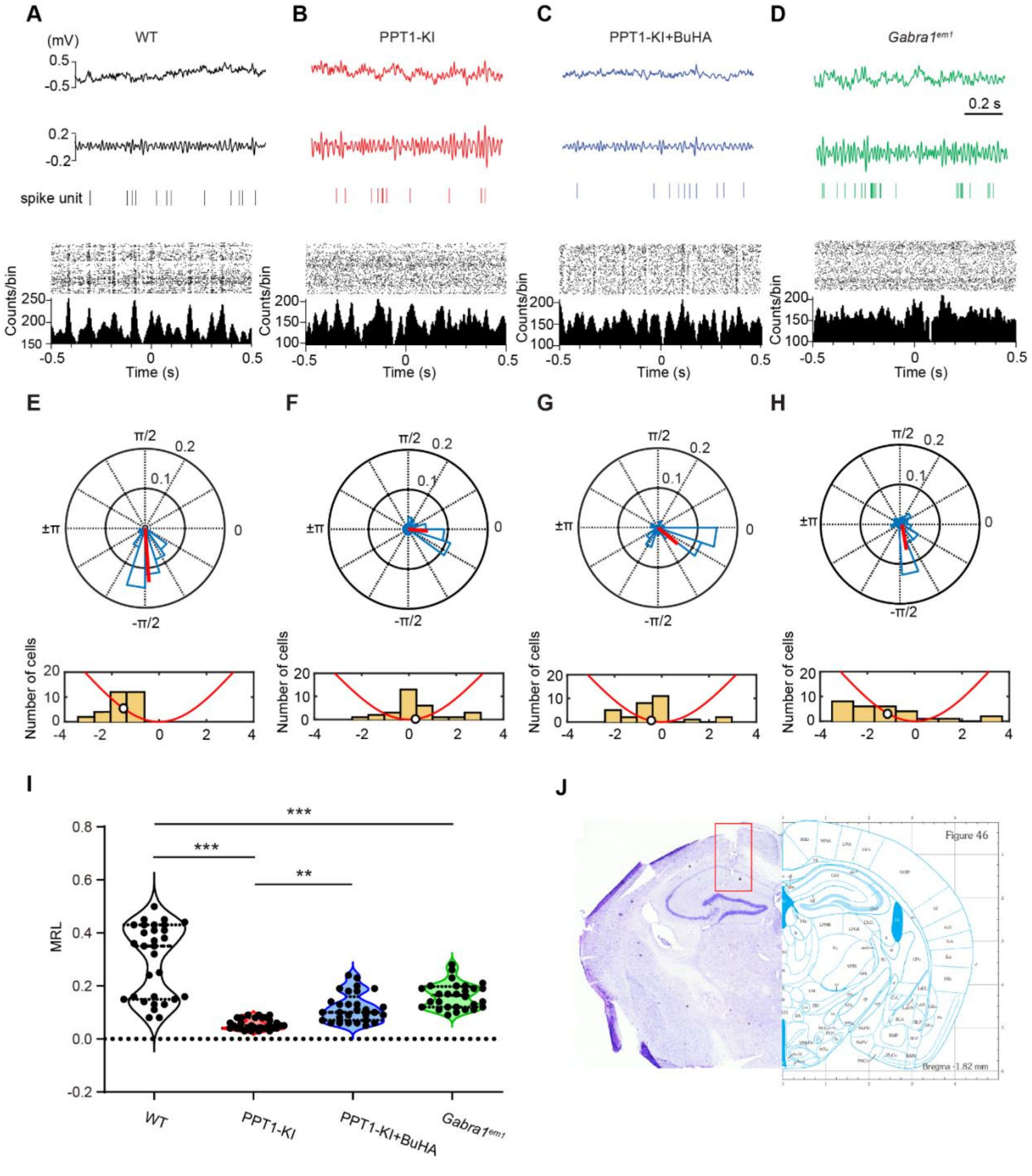


**Figure S8. PPT1 deficiency disrupts theta phase coupling.**

**(A-D)** Sample traces showing gamma phase locking recorded at CA1 region. Peri-event raster (the upper panel) and histogram (the lower panel) displaying phase-coupling of spike units and gamma waves recorded from WT (**A**), PPT1-KI (**B**), BuHA treated PPT1-KI (**C**), and *Gabra1*$^{em1}$ (**D**) mice.

The tips displaying timestamps of spike units. The dotted lines point out the valley of gamma wave. Bin width is 2 ms.

**(E-H)** Circular distribution of the mean-spike gamma-phase angles (15°bin width) (the upper panel) recorded from CA1 area of WT **(E)**, PPT1-KI **(F)**, BuHA treated PPT1-KI **(G)**, and *Gabra1$^{em1}$* **(H)** mice. Red bar displaying the direction and magnitude (length) of the MRL for the population. Distribution of mean-spike gamma-phase angles (45°bin width) (the lower panel). The red line displaying one schematic gamma cycle, the white circle represented the mean phase angle.

**(I)** Comparison of mean MRL value between multi groups. WT: n=30 neurons from 7 mice; PPT1-KI: n= 30 neurons from 7 mice; BuHA treated PPT1-KI: n= 29 neurons from 6 mice; *Gabra1$^{em1}$*: n= 28 neurons from 6 mice. Kruskal-Wallis test, $^{**}P< 0.01$, $^{***}P< 0.001$.

**(J)** Nissl stain indicating the electrode track.

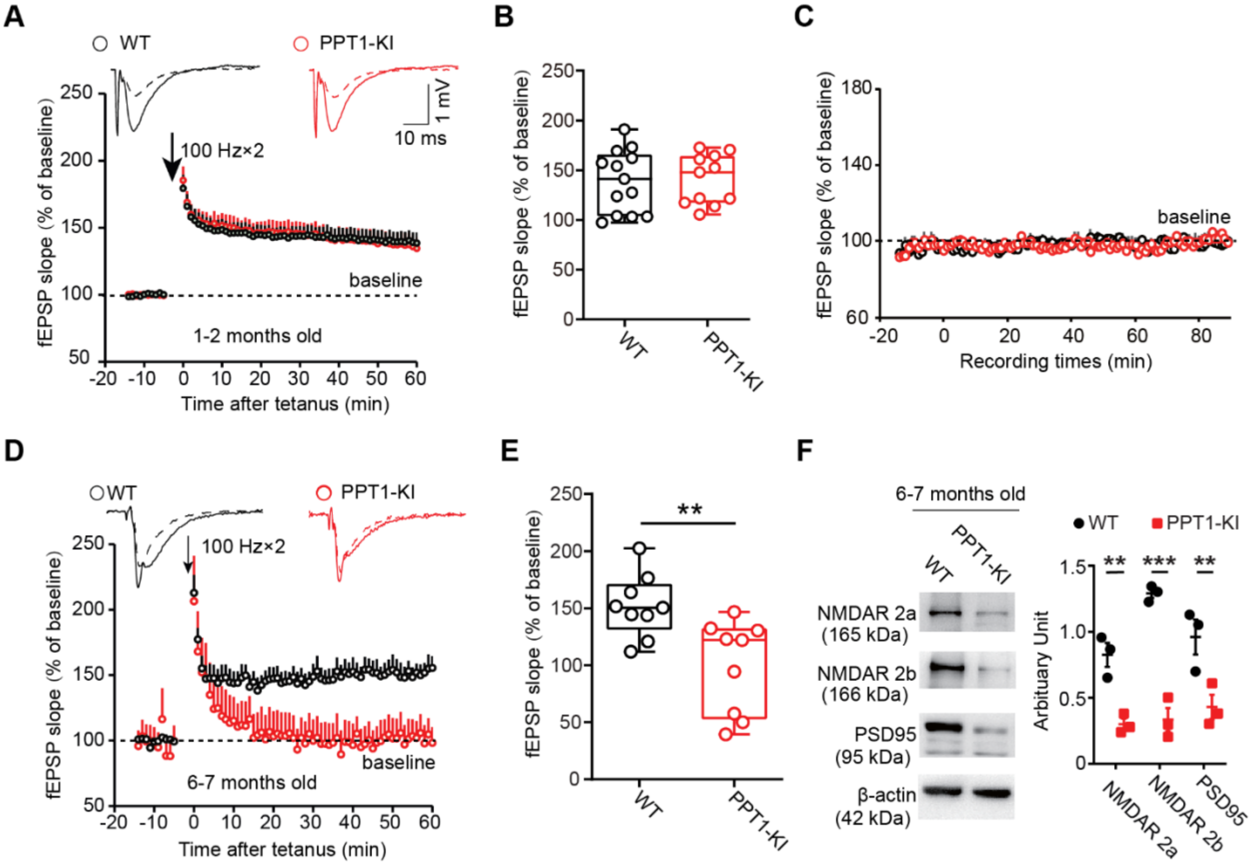


**Figure S9. LTP is impaired in old but not young PPT1-KI mice.**

**(A, B)** Scatter plots showed the nornalized median fEPSP slope before and after tetanus stimulated at SC-CA1 synapse from 1-2 months old mice. The black circles indicated the WT group. The red squares indicated the PPT1-KI groups. Representative traces 5 min before (dotted line) and 50 min after (full line) induction of LTP were shown above the scatter plot **(A)**. Box plots displayed the values of fEPSP slopes between 40-60 min after tetanus **(B)**. WT mice: n=11 slices from 10 mice; PPT1-KI mice: n=13 slices from 11 mice, t-test, no significant difference.

**(C)** Scatter plots showed the nornalized median fEPSP slope without tetanus at SC-CA1 synapse.WT mice: n= 7 slices from 5 mice; PPT1-KI mice: n= 6 slices from 4 mice.

**(D, E)** Scatter plots showed the nornalized median fEPSP slope before and after tetanus stimulated at SC-CA1 synapse from 6-7 months old mice. The black circles indicated the WT group. The red squares indicated the PPT1-KI groups. Representative traces 5 min before (dotted line) and 50 min after (full line) induction of LTP were shown above the scatter plot **(D)**. Box plots displayed the values of fEPSP slopes between 40-60 min after tetanus **(E)**. WT mice: n= 9 slices from 8 mice; PPT1-KI mice: n= 9 slices from 7 mice, t-test, no significant difference, t-test, $^{**}P < 0.01$.

**(F)** Representative immunoblot (left) and quantification of NMDAR 2a/b and PSD95 (right) levels in WT and PPT1-KI slices at 6 to 7 months old mice. N=3 for each group, t-test, $^{**}P < 0.01$, $^{***}P < 0.001$. Data are represented as mean ± SEM.

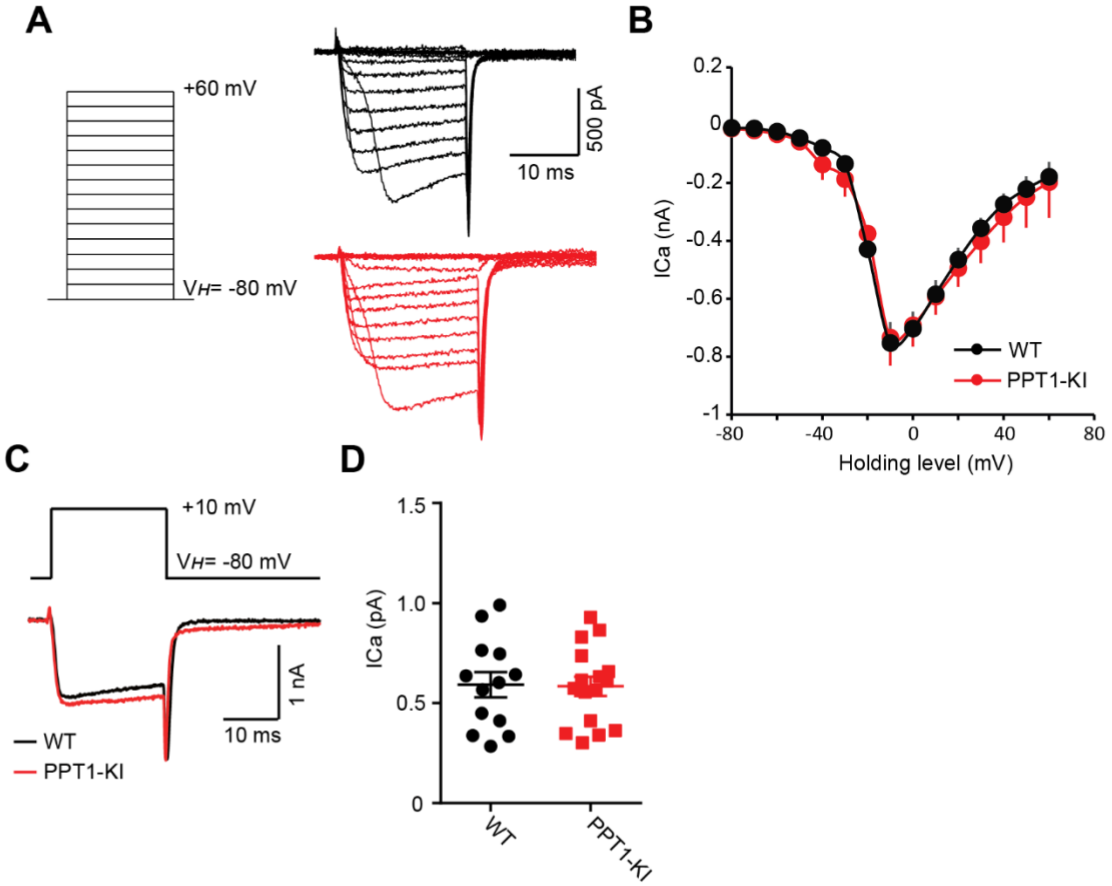


**Figure S10. Voltage-gated calcium channel activation is not disturbed in 1 to 2 months old PPT1-KI mice.**

**(A)** Sample traces of voltage-gated calcium currents evoked by stimulus epochs (-80 to +60 mV by 10 mV steps at 20 ms duration) recorded from WT (black trace) and PPT1-KI (red trace) CA1 pyramidal cells.

**(B)** Curve plotting of current-voltage relationship recorded from WT (black curve, n=10 cells from 8 slices and 8 mice) and PPT1-KI (red curve, n=13 cells from 9 slices and 6 mice) CA1 pyramidal cells. Two-way ANOVA, no significant difference.

**(C)** Sample traces of voltage-gated calcium currents evoked from WT (black trace) and PPT1-KI (red trace) CA1 pyramidal cells by the depolarization from -80 mV to +10 mV at 20 ms duration.

**(D)** Statistical results displaying peak amplitudes of calcium current. WT mice: n=13 cells from 8 slices and 8 mice; PPT1-KI mice: n=16 cells from 9 slices and 6 mice, t-test, no significant difference. Data are represented as mean ± SEM.

### Reference


Morris, R. (1984). Developments of a water-maze procedure for studying spatial learning in the rat. Journal of neuroscience methods *11*, 47-60.

Pietropaolo, S., Sun, Y., Li, R., Brana, C., Feldon, J., and Yee, B.K. (2009). Limited impact of social isolation on Alzheimer-like symptoms in a triple transgenic mouse model. Behavioral neuroscience *123*, 181-195. 10.1037/a0013607.

Vorhees, C.V., and Williams, M.T. (2006). Morris water maze: procedures for assessing spatial and related forms of learning and memory. Nature protocols *1*, 848-858. 10.1038/nprot.2006.116.
